# Ratiometric bioluminescent sensors towards *in vivo* imaging of bacterial signaling

**DOI:** 10.1101/798140

**Authors:** A. B Dippel, W. A. Anderson, J. H. Park, F. H. Yildiz, M.C. Hammond

## Abstract

Second messenger signaling networks allow cells to sense and adapt to changing environmental conditions. In bacteria, the nearly ubiquitous second messenger molecule cyclic di-GMP coordinates diverse processes such as motility, biofilm formation, and virulence. In bacterial pathogens, these signaling networks allow the bacteria to survive changing environment conditions that are experienced during infection of a mammalian host. While studies have examined the effects of cyclic di-GMP levels on virulence in these pathogens, it has previously not been possible to visualize cyclic di-GMP levels in real time during the stages of host infection. Towards this goal, we generate the first ratiometric, chemiluminescent biosensor scaffold that selectively responds to c-di-GMP. By engineering the biosensor scaffold, a suite of Venus-YcgR-NLuc (VYN) biosensors is generated that provide extremely high sensitivity (K_D_ < 300 pM) and large BRET signal changes (up to 109%). As a proof-of-concept that VYN biosensors can image cyclic di-GMP during host infection, we show that the VYN biosensors function in the context of a tissue phantom model, with only ∼10^3^-10^4^ biosensor-expressing cells required for the measurement. Furthermore, the stable BRET signal suggests that the sensors could be used for long-term imaging of cyclic di-GMP dynamics during host infection. The VYN sensors developed here can serve as robust *in vitro* diagnostic tools for high throughput screening, as well as genetically encodable tools for monitoring the dynamics of c-di-GMP in live cells, and lay the groundwork for live cell imaging of c-di-GMP dynamics in bacteria during host infection, and other complex environments.

## INTRODUCTION

The second messenger molecule cyclic di-GMP (c-di-GMP) is a key regulator of bacterial physiology and behavior, coordinating diverse processes such as motility, biofilm formation, and virulence. First discovered as a stimulator of cellulose synthesis,^1^ c-di-GMP has since been found to be nearly ubiquitous in bacteria, with c-di-GMP signaling pathways often integrated with other global regulatory systems, such as phosphorylation networks and quorum sensing pathways.^2,3^ The intracellular levels of c-di-GMP are tightly regulated by diguanylate cyclase (DGC) and phosphodiesterase (PDE) enzymes that synthesize and degrade c-di-GMP, respectively. Many bacteria have an abundance of predicted DGC and PDE genes, suggesting unique c-di-GMP regulatory circuits are activated in response to different environmental cues. In many bacterial pathogens, including *Pseudomonas aeruginosa, Clostridium difficile, Vibrio cholerae*, and pathogenic strains of *Escherichia coli*, these complex c-di-GMP signaling networks allow the bacteria to adapt to and survive in the changing environmental conditions that are experienced during infection of a mammalian host.^4^ While multiple studies have examined the effects of c-di-GMP levels on virulence in these pathogens, there currently are no tools available that allow for the quantification of c-di-GMP in bacteria during stages of mammalian host infection. To interrogate these complex c-di-GMP signaling networks in bacteria over the course of the infection process, new analytical tools are needed for quantifying and imaging intracellular c-di-GMP levels within tissue over extended time frames.

Commonly used tools for analyzing intracellular c-di-GMP levels include phenotypic screens and mass spectrometry (MS) analysis of bacterial cell extracts. Phenotypic screens for motility and biofilm formation can serve as proxies for measuring intracellular c-di-GMP levels.^5,6^ These assays can be high throughput and are useful in screening genetic knockouts, however they have low sensitivity and provide indirect measurement of c-di-GMP that can be complicated by pleiotropic effects. MS-based analysis of c-di-GMP from bacterial cell extracts is highly sensitive and quantitative, however the multi-step sample preparation and long analysis time required leads to reduced throughput.^7–10^ In addition, neither phenotypic assays nor mass spectrometry-based analysis can provide real-time, dynamic measurements of c-di-GMP in cells. To overcome these issues, our lab and others have developed several genetically encodable fluorescent biosensors that can report on single-cell dynamics of c-di-GMP using fluorescence microscopy or flow cytometry.^11–13^ These tools are sensitive, can provide real-time measurements of c-di-GMP dynamics, and are amenable to high throughput screening. Notable examples include protein-based FRET biosensors that have been used to image c-di-GMP dynamics during asymmetric cell division in *Caulobacter crescentus*^11^ and RNA-based fluorescent sensors that were used to visualize c-di-GMP changes in *E. coli* in direct response to an environmental signal, zinc.^14^

One drawback of fluorescent biosensors, however, is that due to a reliance on external illumination, these systems are incompatible with imaging in deep tissues of animals and in long-term experiments due to phototoxicity and/or photobleaching. In a preliminary effort to expand the capabilities of genetically encodable tools for quantifying c-di-GMP levels to overcome these issues, our lab developed the first chemiluminescent biosensors for c-di-GMP^15^ based on the yellow Nano-lantern (YNL) scaffold and a CSL-BRET mechanism.^16^ These YNL-YcgR biosensors provide nanomolar sensitivity for c-di-GMP with high selectivity, large signal changes, and a luminescent signal that is produced without external illumination. The sensors were used to develop a rapid, plate-reader based assay for measuring diguanylate cyclase activity in bacterial lysates. The intensity-based signal of these sensors is useful for *in vitro* activity assays with lysates or purified enzymes where biosensor and luminescent substrate levels can be controlled. However, in long-term imaging experiments and/or situations where luminescent substrate availability differs between samples, signal quantitation becomes complicated for intensity-based sensors. Accordingly, we encountered issues when applying the YNL sensors to live cell measurements of c-di-GMP in bacteria.^15^

Ratiometric BRET sensors using the engineered marine luciferase NanoLuc (NLuc)^17^ have recently been developed for imaging Ca^2+^,^18^ Zn^2+^,^19^ and membrane voltage,^20^ however to our knowledge no sensors of this type have been applied to imaging in bacteria to date. In this work, we generate a suite of BRET biosensors that selectively respond to c-di-GMP and produce ratiometric signal changes. The tVYN-TmΔ biosensor was applied in a plate reader-based assay to quantify c-di-GMP levels in *V. cholerae* extracts grown under a variety of conditions that mimic the infection cycle, with quantitation and sensitivity comparable to MS-based methods (LOD = 30 fmol). We also show that luminescent biosensors producing ratiometric-based signal enable long-term, quantitative, live cell imaging of c-di-GMP activity in *E. coli* using an IVIS small animal imaging system.

## RESULTS AND DISCUSSION

### Design of BRET sensor for cyclic di-GMP

The starting BRET scaffold, V-NLuc, pairs the newly developed marine luciferase, NanoLuc (NLuc), with a truncation of the monomeric yellow fluorescent protein, Venus, as the donor and acceptor moieties, respectively, similar to the previously developed YeNL.^21^ NLuc produces a glow-type luminescence with an emission maximum at 460 nm and an overall luminescent output ∼100-150x that of the commonly used Renilla or firefly luciferases.^17^ Compared to the intensity-based yellow Nano-lantern (YNL) sensors for c-di-GMP previously designed by our lab that use a mutated version of Renilla luciferase as the donor moiety, the substitution of NLuc should produce significantly higher signal intensity, improved thermodynamic stability, and increased signal stability over time. The emission of NLuc overlaps well with the excitation of Venus, producing an efficient energy transfer in V-NLuc, as measured by the BRET ratio (530/460 nm) (Figure 1a).

**Figure 1.**
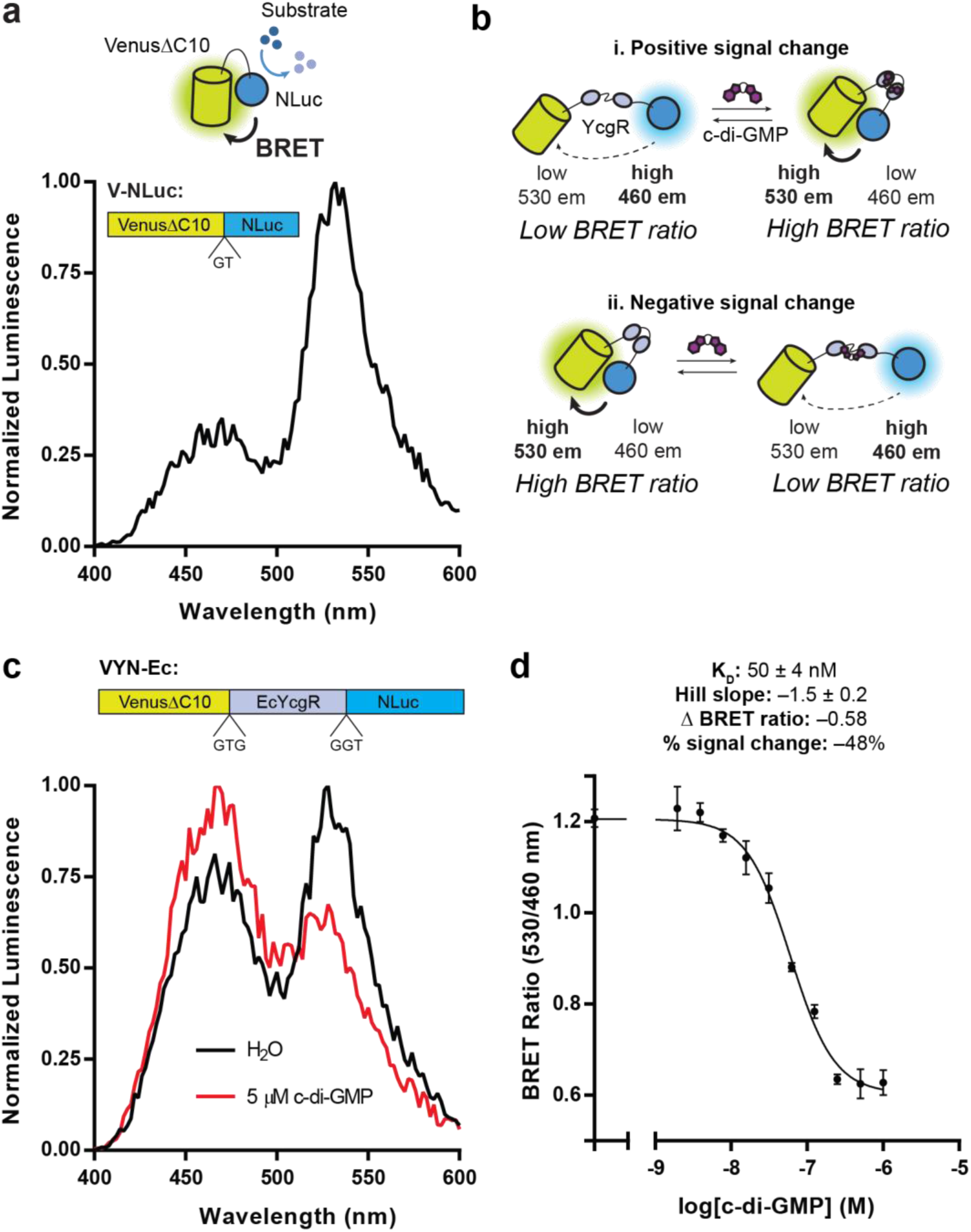
Design and characterization of ratiometric VYN biosensors. (a) Schematic for BRET mechanism of V-NLuc scaffold and the domain structure of the protein. Normalized luminescence emission spectra of purified V-NLuc. Data from one representative measurement. (b) Two potential mechanisms for modulation of BRET ratios by c-di-GMP binding to VYN sensors. (c) Normalized luminescence emission spectra of purified VYN-Ec in the presence and absence of c-di-GMP. Data from one representative measurement shows the biosensor follows mechanism ii. Schematic of VYN-Ec sensor shown above. (d) Binding affinity measurements for purified VYN-Ec. Data are from 3 replicates represented as mean +/- SD.

To design a c-di-GMP sensor, a c-di-GMP-binding YcgR protein is inserted between Venus and NLuc (Figure 1b). YcgR-like proteins contain the c-di-GMP-binding PilZ domain at their C-terminus.^22–24^ These proteins typically undergo large conformational changes upon c-di-GMP binding,^24^ which has been harnessed for the generation of genetically encoded sensors for c-di-GMP.^11,12,15,25^ Thus, binding of c-di-GMP to a <u>V</u>enus-<u>Y</u>cgR-<u>N</u>Luc (VYN) sensor should produce a change in energy transfer efficiency between Venus and NLuc, changing the BRET ratio. The change could be from low to high BRET ratio upon binding, or vice versa, thereby producing positive or negative signal changes, respectively (Figure 1b).

For initial testing of the sensor design, full-length YcgR protein from *Escherichia coli* (*Ec*YcgR) was inserted into the BRET scaffold to generate VYN-Ec. The sensor was purified from *E. coli* after co-expression with the c-di-GMP-specific phosphodiesterase (PDE) PdeH to ensure the sensor did not co-purify with endogenous c-di-GMP. The purified sensor showed a c-di-GMP dependent change from high to low BRET state (Figure 1c). Promisingly, the BRET ratio of VYN-Ec remained stable over time even as overall signal intensity decreased due to consumption of luminescent substrate (Figure S1a, b). VYN-Ec binds to two molecules of c-di-GMP with an apparent affinity (KD) of ∼50 nM and a BRET signal change of −48% (Figure 1d). The sensor retains selectivity for c-di-GMP over structurally related cyclic dinucleotides (Figure S1c, d).

The affinity of the VYN-Ec sensor (K_D_∼50 nM) is significantly higher compared to the equivalent YNL-*Ec*YcgR sensor (K_D_∼350 nM)^15^ and previously reported affinity values for *Ec*YcgR (K_D_∼800 nM).^22^ Since c-di-GMP binding to PilZ domains has been found to be largely entropically driven,^24^ this finding suggests that the VYN scaffold itself may be providing a degree of additional stability that results in a decreased entropic cost of binding c-di-GMP, leading to increased binding affinity. Encouraged by the VYN-Ec results, we sought to further improve the properties of the sensor via phylogenetic screening and semi-rational protein engineering.

### Optimization of VYN biosensors

Four phylogenetic YcgR variants previously characterized in the YNL scaffold were selected for testing in the VYN scaffold. These YcgR variants were chosen because in the YNL scaffold they displayed high affinity, high stability, and large positive signal changes (*Tm*YcgR, *Cp*YcgR, and *Tb*YcgR), or a moderate affinity and a moderate negative signal change (*Nt*YcgR).^15^ Purified VYN-Tm, VYN-Nt, and VYN-Tb sensors exhibited c-di-GMP dependent changes in BRET, while the VYN-Cp sensor appeared non-responsive (Figure 2a). As expected, the functional sensors displayed higher affinities for c-di-GMP compared to VYN-Ec.

**Figure 2.**
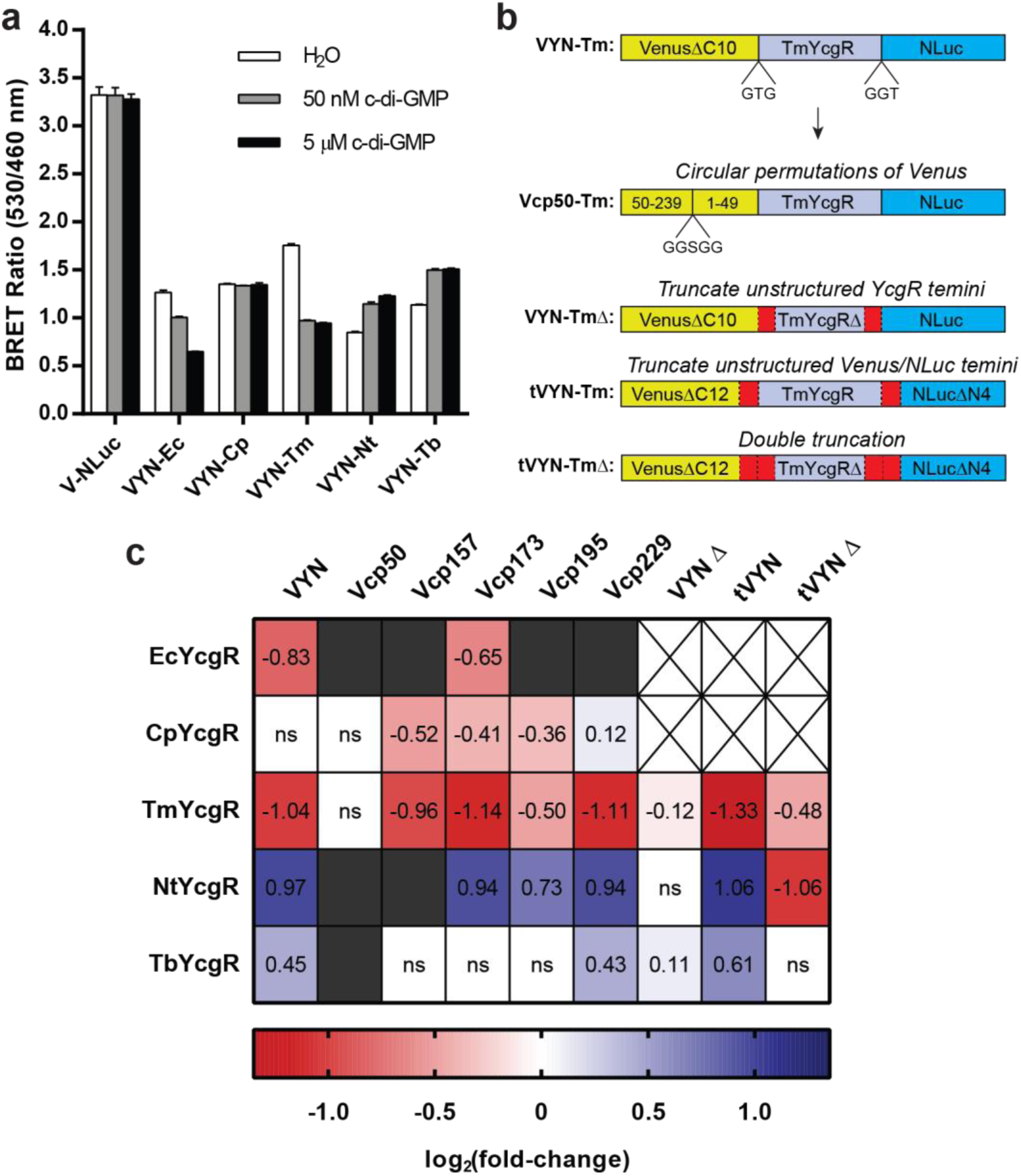
Optimization of VYN sensors for c-di-GMP. (a) BRET ratios of purified VYN sensors containing phylogenetic variant YcgR proteins in response to varying levels of c-di-GMP. Data are from 3 replicates represented as mean +/- SD. (b) Schematic representations of the domain architectures of altered VYN scaffolds, using TmYcgR as an example. Red regions highlight where truncations were made. (c) Signal fold-change (defined as BRET ratio with 5 µM c-di-GMP added divided by BRET ratio with buffer added and plotted as log2(fold-change)) of VYN sensor library screened in lysates. Grayed boxes = dim signal; crossed-out boxes = not tested; ns = no significant signal fold-change (P > 0.05 determined by Student’s t-test). Data are from 4 biological replicates represented as the mean.

We sought to further improve the signal change of these VYN sensors through two routes: composite linker truncation and circular permutation of Venus. Both strategies are commonly used to improve signal change in the development of ratiometric sensors (FRET and BRET), but it is difficult to predict the effects on the resulting sensor.^26,27^ Accordingly, a small library of VYN variants was generated to screen for sensors with improved properties (Figure 2b). For linker truncation, the “composite linkers” (defined as the N- and C-terminal residues of Venus, YcgR, and NLuc that are not necessary for fluorescence, ligand binding, or luminescence) were analyzed. For the truncated VYN (tVYN) scaffold, 2 additional C-terminal residues from Venus and 4 N-terminal residues from NLuc were removed.^21^ For YcgRΔ variants, the secondary structure prediction software SABLE was used to predict unstructured N- and C-terminal residues for removal (Table S2). For circular permutations of Venus, five variants were chosen that were shown to not disrupt Venus fluorescence (Table S3).^28^

A library of 39 VYN variants was constructed and screened in a lysate-based assay that allows for biosensor performance to be rapidly assessed without protein purification. In this assay each individual sensor is co-expressed in *E. coli* with PdeH, cells are lysed, c-di-GMP is added to the lysate at specified concentrations (0, 50 nM, and 5 µM), and BRET ratios are measured. In Figure 2c, the log_2_(signal fold-change) values for each sensor are reported to simplify the comparison of positive and negative signal change sensors. While no clear trends could be drawn between designs, the small set of linker truncations and circular permutations tested produced at least one BRET sensor for each YcgR protein with a signal change of −30% or +50% (Figure S2). Interestingly, this screen showed that seemingly small alterations in the scaffold can produce large differences in signal fold-change. The switch from tVYN-Nt to tVYN-NtΔ, for example, produced a sensor with the same relative signal fold-change, but flipped from a positive to negative change in BRET ratios (Figure 1b).

A subset of the sensors from the library was purified and characterized *in vitro* and was shown to span a range of affinities from < 300 pM up to ∼100 nM (Table 1). The tVYN-TmΔ sensor, to our knowledge, exhibits the highest affinity cyclic dinucleotide:protein interaction ever measured, and the largest magnitude signal change (Δ ratio of –1.04) out of all tested sensors *in vitro*. Interestingly, this sensor exhibited larger signal changes *in vitro* than in lysates, mostly due to truncated NLuc protein obscuring the BRET ratio in unpurified lysates (Figure S3). Given its desirable properties, we chose to apply the tVYN-TmΔ sensor to develop a plate reader assay for quantification of c-di-GMP in cell extracts.

**Table 1.** Characteristics of selected VYN sensor variants. Notes: ^a^Data are from 3 replicates represented as mean ± SD. ^b^Affinity measurements were made using 300 pM biosensor to determine K_D_ values <3 nM. ^c^Biosensor constructs were purified using an N-terminal His_6_ tag, as opposed to a C-terminal His_6_ tag for all others.

| Sensor | $\Delta$ ratio<sup>a</sup> | % change<sup>a</sup> | $K_D$ (nM)<sup>a</sup> | Hill coefficient<sup>a</sup> |
| --- | --- | --- | --- | --- |
| tVYN-TmΔ<sup>b, c</sup> | -1.04 | -56% | $<0.3$ | $-1.7 \pm 0.03$ |
| tVYN-Tm<sup>b</sup> | -0.47 | -50% | $0.8 \pm 0.03$ | $-1.7 \pm 0.1$ |
| VYN-Tm<sup>b, c</sup> | -0.67 | -42% | $0.8 \pm 0.2$ | $-1.5 \pm 0.4$ |
| Vcp229-Tm<sup>b</sup> | -0.41 | -44% | $2.0 \pm 0.1$ | $-1.5 \pm 0.1$ |
| VYN-Tb<sup>c</sup> | 0.38 | 33% | $8.0 \pm 0.4$ | $2.4 \pm 0.3$ |
| tVYN-Tb | 0.18 | 38% | $12 \pm 1$ | $2.0 \pm 0.3$ |
| Vcp229-Nt | 0.22 | 52% | $14 \pm 1$ | $1.6 \pm 0.1$ |
| VYN-Nt<sup>c</sup> | 0.39 | 45% | $14 \pm 4$ | $1.4 \pm 0.4$ |
| Vcp173-Nt | 0.04 | 17% | $17 \pm 4$ | $2.0 \pm 0.7$ |
| tVYN-Nt | 0.24 | 56% | $20 \pm 1$ | $1.6 \pm 0.2$ |
| VYN-Ec<sup>c</sup> | -0.58 | -48% | $50 \pm 4$ | $-1.5 \pm 0.2$ |
| tVYN-NtΔ | -1.01 | -51% | $54 \pm 4$ | $-1.7 \pm 0.1$ |
| Vcp157-Cp | -0.39 | -33% | $96 \pm 6$ | $-1.9 \pm 0.2$ |

### Quantification of c-di-GMP in *Vibrio cholerae* cell extracts

The quantification of intracellular c-di-GMP levels is routinely performed using mass spectrometry (MS)-based analysis of bacterial cell extracts. These methods are highly sensitive and allow for the quantitation of c-di-GMP in the picomolar or femtomolar range, depending on the detection method used.^7–9^ However, the sample preparation steps, long analysis time, and expertise required to perform MS-based analysis of cell extracts has limited the accessibility and throughput of these types of experiments.

Given the extremely high affinity of the tVYN-TmΔ sensor for c-di-GMP, we predicted that it would be possible to develop a simple and robust plate reader-based assay for quantification of c-di-GMP. While the sensitivity was highest using 300 pM biosensor (Figure 3a), quantitation of extracts was performed with 3 nM biosensor due to improved signal intensity and stability. Under these conditions, the limit of detection (signal-to-noise ratio 3:1) of the tVYN-TmΔ sensor was measured to be 30 fmol, which is comparable to the most sensitive established LC-MS/MS-based methods (Figure 3b).^7,8^ One drawback of the biosensor assay is the limited linear range (∼30 fmol to 400 fmol), however this can be alleviated by diluting any samples that fall outside of this range (generally by 1:10-1:20) or using more biosensor.

**Figure 3.**
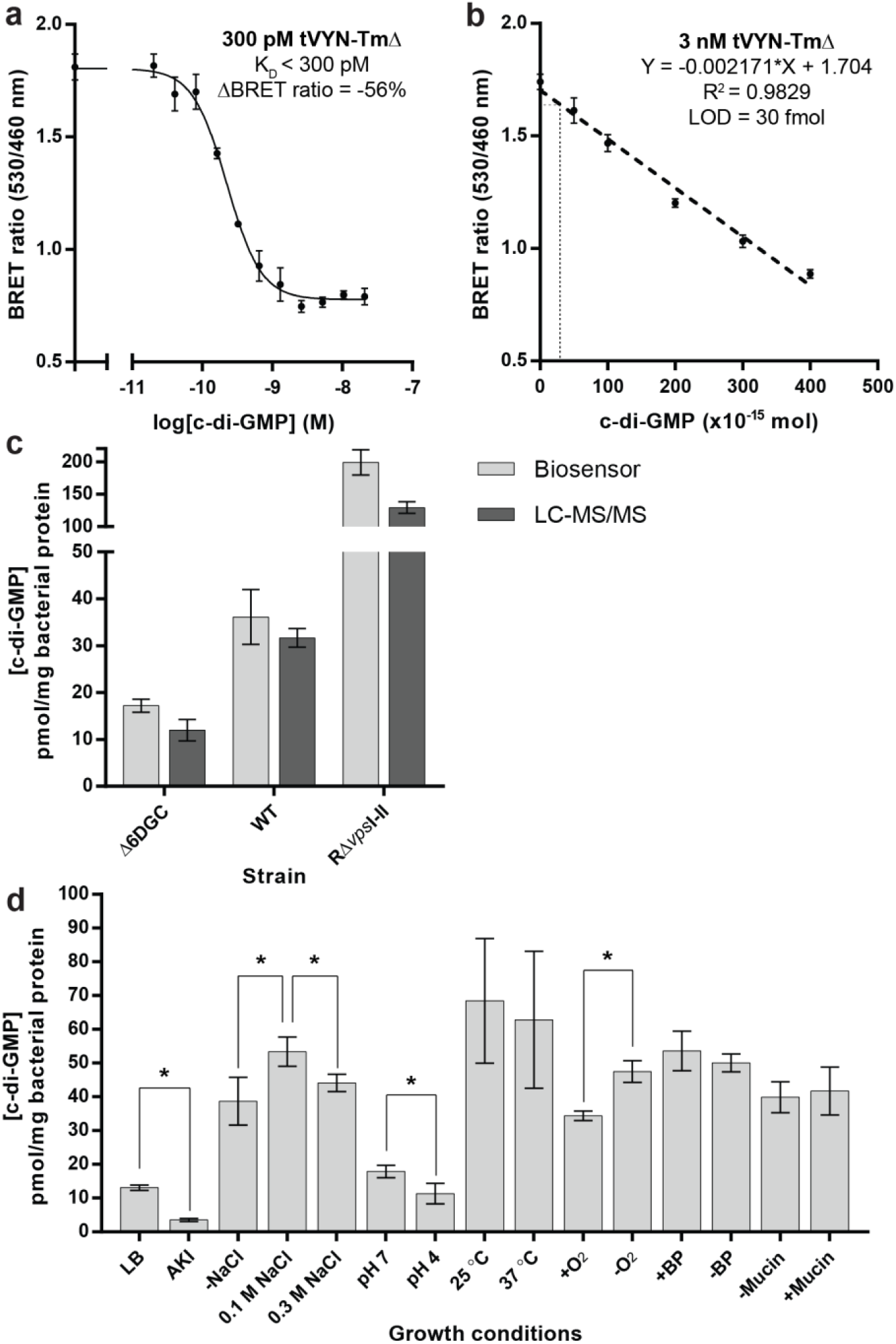
Quantitation of c-di-GMP using tVYN-TmΔ biosensor. (a) Binding affinity measurements for purified tVYN-TmΔ using decreased biosensor concentration. Data are from 3 replicates represented as mean +/- SD. (b) Representative standard curve for c-di-GMP quantitation using purified tVYN-TmΔ. Data are from 6 replicates represented as mean +/- SD. (c) Quantitation of c-di-GMP in cell extracts of 3 different strains of *V. cholerae* using the tVYN- TmΔ biosensor and LC-MS/MS. Data are from 3 biological replicates represented as the mean +/- SD. (d) Quantitation of c-di-GMP in cell extracts of WT *V. cholerae* grown under different conditions using the tVYN-TmΔ biosensor. Asterisks (*) denote significant changes in c-di-GMP between growth conditions (P < 0.05 determined by Student’s t-test). Data are from 3 biological replicates represented as the mean +/- SD, except for 25 and 37 °C conditions, which are from 6 biological replicates.

To directly compare the performance of our plate reader-based protocol to established LC-MS/MS methods, cell extract samples from *V. cholerae* were analyzed using both methods. Cell extracts were generated from three different strains of *V. cholerae* – wild-type (WT), wild-type lacking six DGCs (Δ6DGC), and rugose (RΔvpsI/II) – that were expected to produce endogenous, low, and high levels of c-di-GMP respectively. The expected differences in c-di-GMP were observed between the three strains and the quantitative data were closely correlated between the biosensor and LC-MS/MS measurements (Figure 3c; Figure S4).

We then analyzed c-di-GMP levels for *V. cholerae* grown under a variety of different conditions that have either been shown to affect or are predicted to affect endogenous c-di-GMP signaling networks within the bacterium. Altered c-di-GMP levels were observed as a result of changes in growth conditions such as O_2_ content, virulence factor-inducing media (AKI), pH, and salinity (Figure 3d; Figure S5). We have previously shown that c-di-GMP levels are increased at lower temperature^29^ and while this trend was indeed observed, the differences were not statistically significant due to high variability between biological replicates. No change was observed with iron or mucin.

Increased levels of c-di-GMP in low O_2_ conditions are consistent with the higher DGC activity of the diferrous form of Vc Bhr-DGC that is stable only under anaerobic conditions.^30^ It has been reported that *V. cholerae* cells exponentially grown in AKI did not show significantly different c-di-GMP levels compared to LB-grown cells,^10^ however the cells were harvested before stationary phase in that study, whereas they were harvested after stationary phase in our study. How pH affects c-di-GMP level in *V. cholerae* has not been shown, so further studies are required to explore this mechanism. This study also provides the first evidence that salinity directly affects c-di-GMP levels in *V. cholerae*. The non-linear results are consistent with the increased expression levels of the first gene in the c-di-GMP-regulated *vps* cluster, *vpsL*, at median (0.1 M) salinity compared to low (0 M) and high (0.2-0.5 M) salinity.^31^

The limit of detection of the biosensor assay is comparable to the most sensitive established LC-MS/MS methods, but is plate-based and significantly more rapid, which makes it well-suited to high-throughput screening applications, such as monitoring enzyme activity or high throughput screening of activators/inhibitors of DGCs and PDEs.^32^ Accordingly, we determined the Z’ factor to be >0.5 for the duration of 30 min after substrate addition (Figure S6), which is considered to be excellent statistical reliability for high-throughput screening.^33^ We even have found that the biosensor signal is sufficiently bright to be analyzed using a digital camera, which drastically reduces the cost of hardware required for c-di-GMP quantification. When applied to analysis of *V. cholerae* extracts, the biosensor assay was able to reliably quantify c-di-GMP concentrations for different strains and under different growth conditions. These results suggest that the biosensor assay will be generally applicable to the study of c-di-GMP in complex bacterial extract samples, including clinical isolates and mixed cultures.

### Live-cell measurements of c-di-GMP using IVIS imager and tissue-like phantom model

With our prior YNL-based sensors we encountered difficulties in making live cell measurements, likely due to changes in luminescent substrate availability and biosensor expression between different conditions that complicated normalization of the intensity-based signal (Figure S7).^15^ To test if the ratiometric VYN biosensors alleviate these issues for live cell measurements, a subset of sensors were co-expressed in BL21 Star (DE3) *E. coli* with a PDE (PdeH – low c-di-GMP), an inactive diguanylate cyclase (DGC) as a control (WspR-G249A – endogenous c-di-GMP), or a constitutively active DGC (WspR-D70E – elevated c-di-GMP). Encouragingly, many of the sensors showed significant changes in BRET ratio between low or endogenous c-di-GMP conditions versus elevated c-di-GMP conditions (Figure 4).

**Figure 4.**
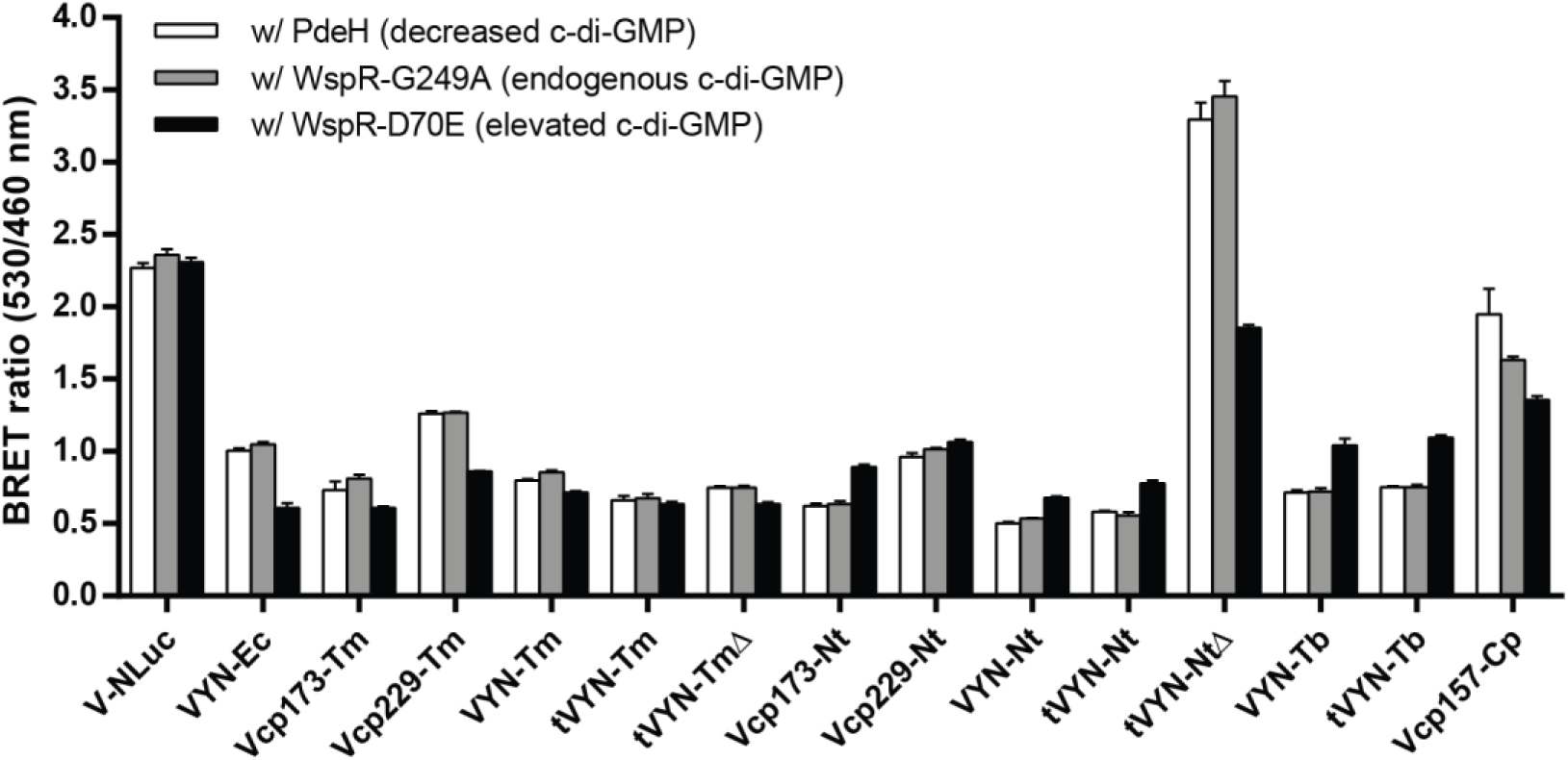
Live cell measurements of c-di-GMP using VYN sensors. BRET ratios of *E. coli* cells co-expressing VYN biosensors with PdeH, WspR-G249A, or WspR-D70E. Data are from 4 biological replicates represented as mean +/- SD.

One of our long-term goals is to monitor signaling activity of bacterial cells in real time during host infection. An initial proof-of-concept experiment was to validate the signal intensity and BRET signal changes of our sensors measured in an instrument routinely used for non-invasive small animal imaging, a Xenogen IVIS 100, with conventional filter sets and settings. Selected sensors were co-expressed in BL21 Star (DE3) *E. coli* cells with WspR-G249A or WspR-D70E to produce endogenous or elevated c-di-GMP levels, respectively. Cells were prepared in the same manner as for plate reader experiments, then images were captured sequentially after luminescent substrate addition using no emission filter and standard 500 and 540 nm emission filters on the IVIS. The total flux (photons/sec) from each well in the 540 nm and 500 nm filter images was used to calculate BRET ratios (Figure 5a).

**Figure 5.**
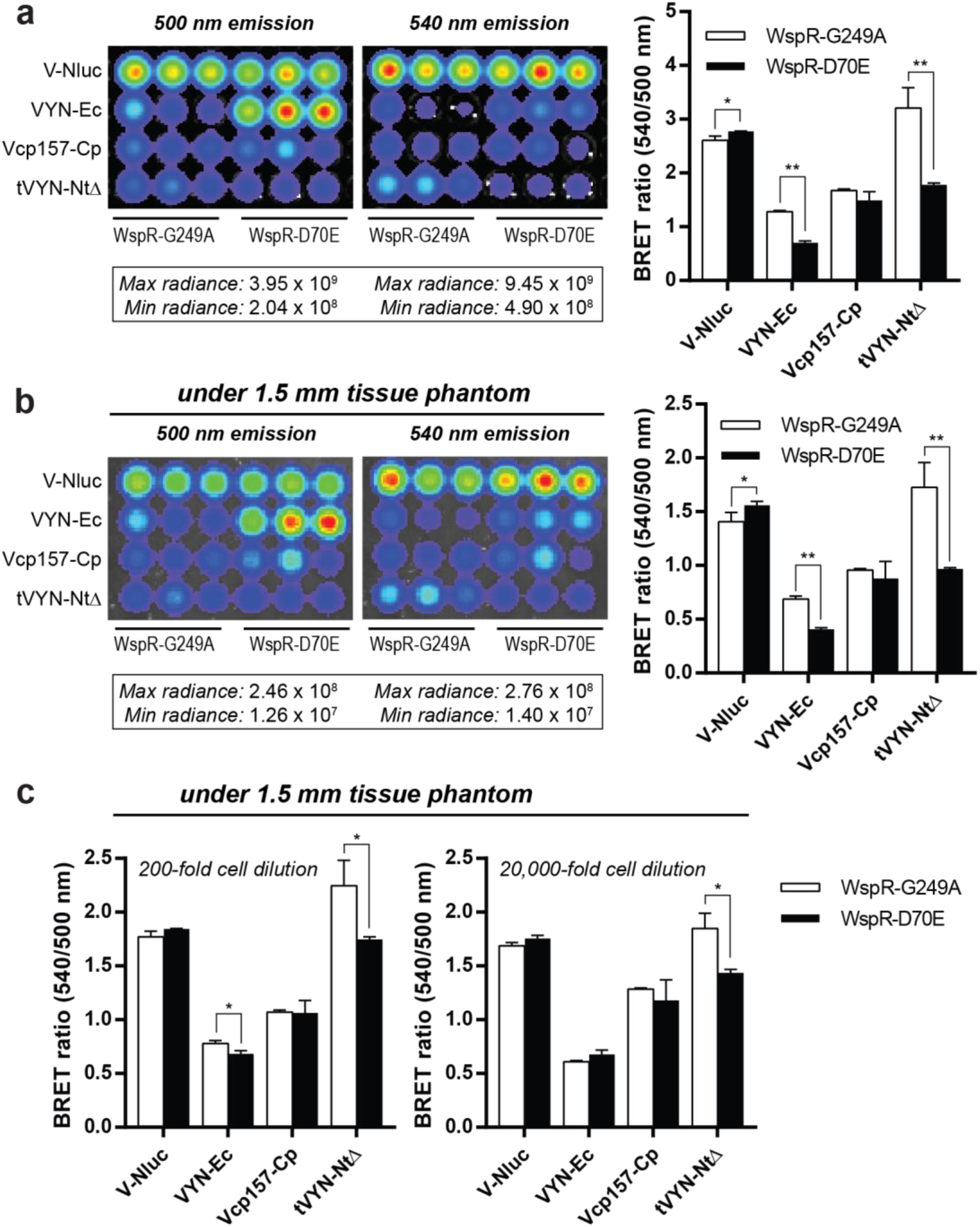
Live cell measurements of c-di-GMP using a tissue phantom model. (a) Luminescent images of 2-fold diluted *E. coli* cultures co-expressing VYN biosensors with WspR-G249A or WspR-D70E captured by an IVIS 100, and the BRET values calculated from the total radiance in each well. Maximum and minimum radiance values (photons/sec/cm^2^ /steradian) captured for each image are shown. (b) Same as part (a), except plate was covered with a 1.5 mm thick tissue phantom prior to image capture. (c) BRET ratios of serially diluted *E. coli* cultures co-expressing VYN biosensors with WspR-G249A or WspR-D70E calculated from the radiance of each well. Plate was covered with a 1.5 mm thick tissue phantom prior to image capture. For all graphs, data are from 3 biological replicates represented as mean +/- SD. Asterisks (*) denote significant changes in BRET ratio (*P < 0.05, **P < 0.005 determined by Student’s t-test).

The raw BRET ratio values are different between IVIS and plate reader instruments, likely due to less optimal emission filters for the biosensor on the IVIS. Nevertheless, the changes in BRET ratio in response to c-di-GMP were faithfully reproduced for VYN-Ec and tVYN-NtΔ sensors with 2- and 200-fold fold dilution of cells (Figures 5a, S8a). With much higher dilution (20,000-fold) and thus lower signal-to-noise, the tVYN-NtΔ sensor but not VYN-Ec maintained the expected response to c-di-GMP, due to its larger magnitude signal change (ΔBRET ∼1 vs. ∼0.5).

To further extend the proof-of-concept, tissue-like phantom materials were utilized to mimic the light absorption and scattering of living tissue.^34^ These types of tissue phantoms have recently been used as a benchmark to compare photon output of luminescent protein systems within deep tissues.^35^ The 96-well plate containing bacterial cells was covered with 1.5 mm thick tissue phantom prior to image capture (Figure S9). While luminescent intensity and BRET ratios were generally lower with application of the tissue phantom (the latter due to hemoglobin absorbing more strongly at 540 nm than 500 nm),^36^ the overall results were similar to without phantom (Figure 5b). Luminescent signal still could be detected down to 20,000-fold dilution for all samples, and the tVYN-NtΔ sensor displayed significant response to c-di-GMP down to that dilution (Figure 5c).

To determine the amount of bacteria monitored in the IVIS experiments, the number of colony-forming units (CFUs) was measured for representative cultures co-expressing the tVYN-NtΔ biosensor and WspR-G249A or WspR-D70E. Cells were prepared as before and then spotted onto LB/Agar plates containing no antibiotic, carbenicillin (Carb), kanamycin (Kan), or both Carb and Kan, the last condition being the overnight growth conditions used for all live-cell experiments. Results from plates with no antibiotics show that there are ∼10^8^ *E. coli* in each well for 2-fold diluted cultures, and ∼10^6^ and ∼10^4^ cells in the 200-fold and 20,000-fold diluted cultures, respectively, as expected. However, comparisons to antibiotic plates reveal that ∼90% of these cells have lost both biosensor and WspR expression plasmids after overnight growth (Figures S10). Thus, the actual number of bacteria producing luminescent signal is only 10% of the total. Given the tVYN-NtΔ sensor is capable of imaging c-di-GMP levels in 20,000-fold diluted cultures in a tissue-like model, this corresponds to as few as ∼10^3^ biosensor-expressing bacterial cells. In comparison, a *V. cholerae* infection model of infant mice presented 10^4^ to 10^5^ bacteria in the small intestine after infection.^37^

Importantly, we found that the BRET ratio signal for tVYN-NtΔ sensor under the tissue phantom remained stable over the course of an hour after luminescent substrate addition, even while overall signal intensity decayed as substrate was consumed (Figure S8c). While the tissue phantom model does not account for substrate distribution *in vivo*, NLuc and furimazine previously have been applied to study the spread of pathogens in mice in real time.^38,39^ Our experiments were performed using coelenterazine-h as the luminescent substrate, so even brighter luminescent signal *in vivo* should be possible with furimazine, which produces higher luminescent output than coelenterazine-h.^17^ Beyond the hour timescale, longer term experiments (days or weeks) could be carried out with repeated administration of luminescent substrate.

## CONCLUSIONS

The work here presents, to our knowledge, the first ratiometric, luminescent biosensors developed to study bacterial signaling. The highest affinity VYN biosensor, tVYN-TmΔ, can serve as an easy-to-use diagnostic reagent for quantifying c-di-GMP levels from bacterial extracts, with comparable sensitivity to LC-MS/MS. Furthermore, as a genetically encodable tool, the tVYN-NtΔ sensor holds considerable promise for monitoring c-di-GMP dynamics in real-time during host infection using a standard small animal imaging system. More broadly, this study demonstrates how to develop and characterize luminescent biosensors that are directed towards studying bacterial activity in complex environments.

## MATERIALS AND METHODS

### Chemiluminescence measurements with purified protein

Briefly, proteins and ligands were prepared in opaque white 96-well LUMITRAC 600 plates (Grenier) in assay buffer [50 mM HEPES (pH 7.2), 100 mM KCl, 10 mM DTT, 0.1% BSA]. Unless otherwise noted, all measurements using purified protein were made using 3 nM sensor in 100 µL total reaction volume, then incubated at 28 °C for at least 10 min to reach binding equilibrium. Chemiluminescent substrate was prepared by diluting coelenterazine-h to 60 µM in reagent buffer [50 mM HEPES (pH 7.2), 100 mM KCl, 300 mM ascorbate], and equilibrating the solution at RT for at least 30 min. Unless otherwise noted, all biosensor measurements were taken at 28 °C in a SpectraMax i3x plate reader (Molecular Devices) after manually adding 20 µL of chemiluminescent substrate. Emission intensities were measured at 460 and 530 nm with 200 ms integration time at 30 s intervals for 10 min after chemiluminescent substrate addition. In general, BRET ratios were calculated using emission values obtained 2 min after substrate addition. For emission spectrum measurements, emission intensities were measured over the range of 400-600 nm in steps of 2 nm.

### Lysate-based assay for biosensor activity

The lysate-based assay was carried out as previously described, with minor modifications.^15^ Single colonies of BL21 Star (DE3) *E. coli* cells co-transformed with pET21-biosensor and pCOLA-PdeH plasmids were resuspended in 500 µL of P-0.5G non-inducing media [0.5% glucose, 25 mM (NH_4_)_2_SO_4_, 50 mM KH_2_PO_4_, 50 mM Na_2_HPO_4_, 1 mM MgSO_4_]^40^ supplemented with 50 µg/mL carbenicillin and 100 µg/mL kanamycin in 2.2 mL 96-well deep-well plates (VWR). Precultures were grown at 37 °C, 340 rpm for 24 h at which point 5 µL of each was used to inoculate 500 µL of ZYP-5052 autoinduction media [25 mM (NH_4_)_2_SO_4_, 50 mM KH_2_PO_4_, 50 mM Na_2_HPO_4_, 1 mM MgSO_4_, 0.5% (v/v) glycerol, 0.05% glucose, 0.2% α-lactose, 1% tryptone, and 0.5% yeast extract]^40^ supplemented with 50 µg/mL carbenicillin and 100 µg/mL kanamycin. Cultures were grown in ZYP-5052 autoinduction media at 37 °C, 340 rpm for 20 h to express the biosensors, then harvested by centrifugation at 4700 rpm for 10 minutes at 4 °C. Lysates were prepared by removing the supernatant media and resuspending cell pellets in 500 µL of screening buffer [50 mM Tris (pH 7.5), 100 mM KCl, 5% glycerol, 2 mM EDTA, 300 μg/mL lysozyme, 1 mM PMSF]. Cells were incubated for 1 h at 4 °C to gently lyse, and total lysates were centrifuged for 40 min at 4700 rpm at 4 °C to generate clarified lysates.

For chemiluminescence measurements, 5 µL of clarified lysate was mixed with 85 µL screening buffer (-lysozyme, -PMSF) and 10 µL of either buffer, 500 nM c-di-GMP, or 50 µM c-di-GMP [in screening buffer (-lysozyme, -PMSF)] in opaque white 96-well LUMITRAC 600 plates (Greiner) to generate final concentrations of 0, 50 nM, or 5 µM c-di-GMP. Chemiluminescence was measured using the same method described for purified protein, except BRET ratios were calculated using emission values obtained 1 min after substrate addition.

### *Vibrio cholerae* strains and growth conditions

*Vibrio cholerae* O1 El Tor A1552 was used as the wild-type strain and two *V. cholerae* strains, Δ6DGC^29^ and RΔ*vps*I-II,^41^ were used as reference strains with low and high cellular c-di-GMP level, respectively. Strains were grown in Luria–Bertani (LB) medium [1% tryptone, 0.5% yeast extract, 0.2 M NaCl; pH 7.5] with constant shaking at 200 rpm at 37°C unless otherwise indicated. To test the effects of salt concentration, LB supplemented with different concentrations of NaCl (0, 0.1, and 0.3M) were used.^42^ To test the effects of different growth temperature^29^ and oxygen availability, the diluted cultures were grown at 25 and 37 °C to OD_600_ ∼0.5 or aerobically and anaerobically (in a Vinyl Anaerobic Airlock Chamber, Coy Laboratory Products) to OD_600_ ∼0.3. To test the effects of mucin addition^43^ and iron depletion,^44^ overnight-grown cultures were inoculated in a 1:200 dilution in LB supplemented with different components [0.4% (w/v) of bovine submaxillary gland mucin (Sigma-Aldrich), or 200 µM of 2,2’-dipyridyl (Alfa Aesar), respectively] and grown to OD_600_ ∼0.5. To test virulence-inducing conditions, overnight-grown cultures were inoculated in a 1:200 dilution in LB and in a 1:100 dilution in AKI [1.5% Bacto peptone, 0.4% yeast extract, 0.5% NaCl, 0.3% NaHCO_3_]. LB cultures were grown overnight with shaking at 220 rpm at 37 °C. AKI cultures were grown statically at 37°C for 4 hours followed by shaking at 220 rpm at 37 °C overnight.^45^ To test the effect of acidic conditions, overnight-grown cultures were inoculated in a 1:200 dilution in LB (pH 7), grown to OD_600_ ∼0.5, and centrifuged at 1500 × g for 7 minutes. Cell pellets were adapted by resuspending in LB (pH 5.7) and incubating for 1 hour. Adapted cells were centrifugated and resuspended in LB (pH 4) followed by 1 hour incubation.^46^

### Live cell measurements with biosensor co-expression

Single colonies of BL21 Star (DE3) *E. coli* cells co-transformed with pET21-biosensor and pCOLA-PdeH, pCOLA-WspR-G249A, or pCOLA-WspR-D70E plasmids were resuspended and grown in the same manner as previously described for the lysate-based assay. After growth and induction of expression, cells were centrifuged, supernatant media was removed, and cell pellets were resuspended in 500 µL PBS [137 mM NaCl, 2.7 mM KCl, 10 mM Na_2_HPO_4_, 1.8 mM KH_2_PO_4_ (pH 7.4)]. For chemiluminescence measurements, cells were diluted 2-fold with PBS in an opaque 96-well LUMITRAC 600 plate (Greiner) to a final volume of 100 µL. Chemiluminescent substrate was added and emission intensities were measured in the same way as described for purified protein. BRET ratios were calculated using emission values obtained 5 min after substrate addition.

### Live cell measurements in tissue-like phantom model

Tissue-like phantoms were prepared as described previously.^34^ Briefly, the phantom solution mixture was prepared with 10% gelatin, 170 µM bovine hemoglobin, and 1% intralipid in TBS-azide buffer [50 mM Tris-HCl (pH 7.4), 150 mM NaCl, 0.1% NaN_3_]. Phantoms were poured to the desired thickness of 1.5 mm between glass plates to ensure uniformity, then stored at 4 °C.

Chemiluminescence measurements were carried out in a Xenogen IVIS 100 Bioluminescent Imager at the Huntsman Cancer Institute Center for Quantitative Cancer Imaging. Cells were grown and prepared in the same way as the live cell co-expression experiments but were diluted 2-fold, 200-fold, and 20,000-fold in PBS in opaque black 96-well assay plates (CoStar). To image the plates, 20 µL of chemiluminescent substrate was added to each well and plates were placed in the chamber. Luminescent images were captured sequentially using no filter, a 500 nm filter, and a 540 nm filter within a 13 cm field of view. The instrument was set to auto-adjust settings to ensure maximum signal for each image (exposure time of 0.5-60 s, binning of 1x-16x, f/stop of 1). For experiments with tissue-like phantom model, the wells were covered with a phantom immediately after addition of chemiluminescent substrate, and luminescent images were captured as before. For image analysis, a 12×8 ROI grid was applied to each image and used to calculate the flux (photons/s) for each individual well. For time course images, the same plate was repeatedly imaged for up to an hour after the initial addition of chemiluminescent substrate.

