## Supporting Information for "Ratiometric bioluminescent sensors towards *in vivo* imaging of bacterial signaling"

### FOR

### Table of Contents

**Additional methods** ........ S3-S4
**Figure S1.** Additional characterization of VYN-Ec sensor ........ S5
**Figure S2.** Lysate-based screen of VYN biosensor library ........ S6
**Figure S3.** Additional characterization of tVYN-TmΔ biosensor ........ S7-S8
**Figure S4.** Biosensor analysis of *V. cholerae* reference strains ........ S9
**Figure S5.** Biosensor analysis of *V. cholerae* under different growth conditions ........ S10
**Figure S6.** Determination of Z’ factors for plate reader assay ........ S11
**Figure S7.** Live cell imaging with luminescent biosensors ........ S12
**Figure S8.** Additional data for live cell measurements of c-di-GMP using a tissue phantom ........ S13
**Figure S9.** Tissue phantom imaging setup ........ S14
**Figure S10.** CFUs in live cell measurements of c-di-GMP ........ S15
**Table S1.** Amino acid sequences of biosensor plasmids ........ S16
**Table S2.** Amino acid sequences of YcgR proteins ........ S17
**Table S3.** Amino acid sequences for tVYN and Vcp variants ........ S18
**Table S4.** Oligonucleotides used in this study ........ S19-S21
**References** ........ S22

#### ADDITIONAL METHODS

**General reagents and oligonucleotides.** Cyclic dinucleotides were purchased from Axxora, LLC. Coelenterazine-h was purchased from NanoLight Technologies and stored as ~6.15 mM stocks in EtOH at -80 °C. Oligonucleotides used in molecular cloning were purchased from Elim Biopharmaceuticals or the University of Utah HSC Core facility.

**Molecular cloning.** The pNL(1.1) plasmid was obtained from Promega. Plasmids encoding PdeH, WspR alleles, and phylogenetic YcgR variants were previously available in the lab. The V-NLuc base scaffold was generated in the $pRSET_B$ plasmid using Gibson assembly[1] with a pRSET-VenusΔC10 backbone and a NanoLuc insert amplified from pNL(1.1). The final vector was designed to contain a KpnI cut site between VenusΔC10 and NanoLuc and include an N-terminal His-tag. Base scaffolds were similarly generated in the pET21 plasmid containing a C-terminal His-tag. Biosensor constructs in $pRSET_B$ or pET21 were generated via Gibson assembly using KpnI digested base scaffolds and PCR amplified YcgR inserts. In general, pET24-YNL-YcgR biosensor plasmids previously generated in the lab[2] were used as template for PCR amplification of YcgR sequences. Truncated biosensors (tVYN and YcgRΔ modifications) were created from the full-length scaffolds using 'Round-the-horn' mutagenesis and Gibson assembly, respectively. Venus circular permutations were created from the base V-NLuc scaffold via three-piece Gibson assembly ligations.

**Protein purification.** Biosensors were purified as described previously, with minor modifications.[2] Biosensors were purified from either the $pRSET_B$ or the pET21 plasmid after co-transforming the sensor plasmid and the pCOLA-PdeH plasmid in *E. coli* BL21 Star (DE3) cells (QB3 MacroLab). Transformants were cultured in 2xYT media at 37 ºC to OD ~ 1.0 and protein expression was induced for 20 h at 18 ºC with IPTG (0.1 mM IPTG for $pRSET_B$ plasmids or 0.5 mM IPTG for pET21 plasmids). Lysates were prepared and the biosensors were purified via Ni-NTA affinity chromatography as previously described.[2] Elution fractions were concentrated and dialyzed to storage buffer [50 mM HEPES (pH 7.2), 100 mM KCl, 10% (v/v) glycerol] using Amicon Ultra-15 Centrifugal Filter Units (10K MWCO; Millipore), and then flash frozen in liquid nitrogen and stored at -80 ºC. Final protein concentrations were determined using the absorption of Venus at 515 nm (extinction coefficient = 92200 $M^{-1}$ $cm^{-1}$). All proteins were analyzed via SDS-PAGE to confirm purity.

**Bacterial cell extract analysis.** C-di-GMP extraction was performed as previously described with minor modification.[3] Briefly, 1.5 mL of *V. cholerae* culture was centrifuged at 1500 x g for 7 minutes. Cell pellets were allowed to dry briefly then re-suspended in 1 mL extraction solution [40% acetonitrile, 40% methanol, 20% water], and incubated on ice for 5 minutes. Samples were centrifuged at 16,000 x g for 5 minutes and 900 µL of supernatant was dried under vacuum and lyophilized. Samples were re-suspended in 100 µL of $ddH_2O$ and analyzed using the tVYN-TmΔ biosensor or liquid chromatography-tandem mass spectrometry (LC-MS/MS) to determine intracellular c-di-GMP levels. Each extract analysis was performed with three biological replicates.

For biosensor analysis, the re-suspended extracts were serially diluted with $ddH_2O$ and 20 µL of serially diluted extract was mixed with 80 µL of biosensor and assay buffer in opaque white 96-well LUMITRAC 600 plates (Grenier) to give final concentrations of 3 nM biosensor and 1x assay buffer [50 mM HEPES (pH 7.2), 100 mM KCl, 10 mM DTT, 0.1% BSA]. Samples were incubated at 28 ºC for at least 10 min to reach binding equilibrium and chemiluminescence was measured as described for purified protein. BRET values were calculated using emission values obtained 5 minutes after substrate addition and each biosensor measurement was performed with two technical replicates. Mean BRET values that fell within the linear range of the biosensor (as determined by a standard curve with pure c-di-GMP) were used to calculate the total amount of c-di-GMP in each sample, and c-di-GMP amounts were normalized to total protein content.

For LC-MS/MS analysis, NaCl was added to the resuspended samples to a final concentration of 184 mM. Samples were analyzed via LC-MS/MS on a Thermo-Electron Finnigan LTQ mass spectrometer coupled to a surveyor HPLC (Thermo Scientific). The Synergi Hydro 4u Fusion-RP 80A column (150 mm x 2.00 mm diameter; 4-µm particle size) (Phenomenex) was used for reverse-phase liquid chromatography. Solvent A was 0.1% acetic acid in 10 mM ammonium acetate, solvent B was 0.1% formic acid in methanol. The gradient used was as follows: time (t) = 0–4 minutes, 98% solvent A, 2% solvent B; t = 10–15 minutes, 5% solvent A, 95% solvent B. The injection volume was 20 µL and the flow rate for chromatography was 200 µL/minute.

The amount of c-di-GMP in samples was calculated with a standard curve generated from pure c-di-GMP suspended in 184 mM NaCl. Concentrations used for standard curve generation were 25 nM, 50 nM, 100 nM, 500 nM, and 1 µM. The assay is linear from 25 nM to 1 µM with an $R^2$ of 0.999. C-di-GMP levels were normalized to total protein content in each culture.

To determine total protein content, 1.5 mL from each culture was pelleted, the supernatant was removed, and cells were lysed in 1.25 mL of 2% sodium dodecyl sulfate. Total protein in the samples was estimated with the BCA assay (Pierce) using bovine serum albumin (BSA) as standards.

**Measurement of colony forming units (CFUs).** To determine the total number of bacterial cells in the IVIS experiments, cells co-expressing pET21-tVYN-NtΔ and pCOLA-WspR-G249A or pCOLA-WspR-D70E were grown as described for live cell measurements with biosensor co-expression. After growth in ZYP-5052 autoinduction media, cells were centrifuged, supernatant media was removed, and cell pellets were resuspended in 500 µL PBS. Resuspended cultures were serially diluted ($10^{-4}$, $10^{-5}$, $10^{-6}$, and $10^{-7}$) with PBS and 10 µL of each serial dilution was spotted on LB/Agar plates containing no antibiotics, 50 µg/mL carbenicillin, 50 µg/mL kanamycin, or 50 µg/mL of both carbenicillin and kanamycin. Each serial dilution was spotted in duplicate, and two biological replicates were used for each culture. Plates were incubated at 37 °C overnight, and colonies were counted to determine total CFUs/mL.

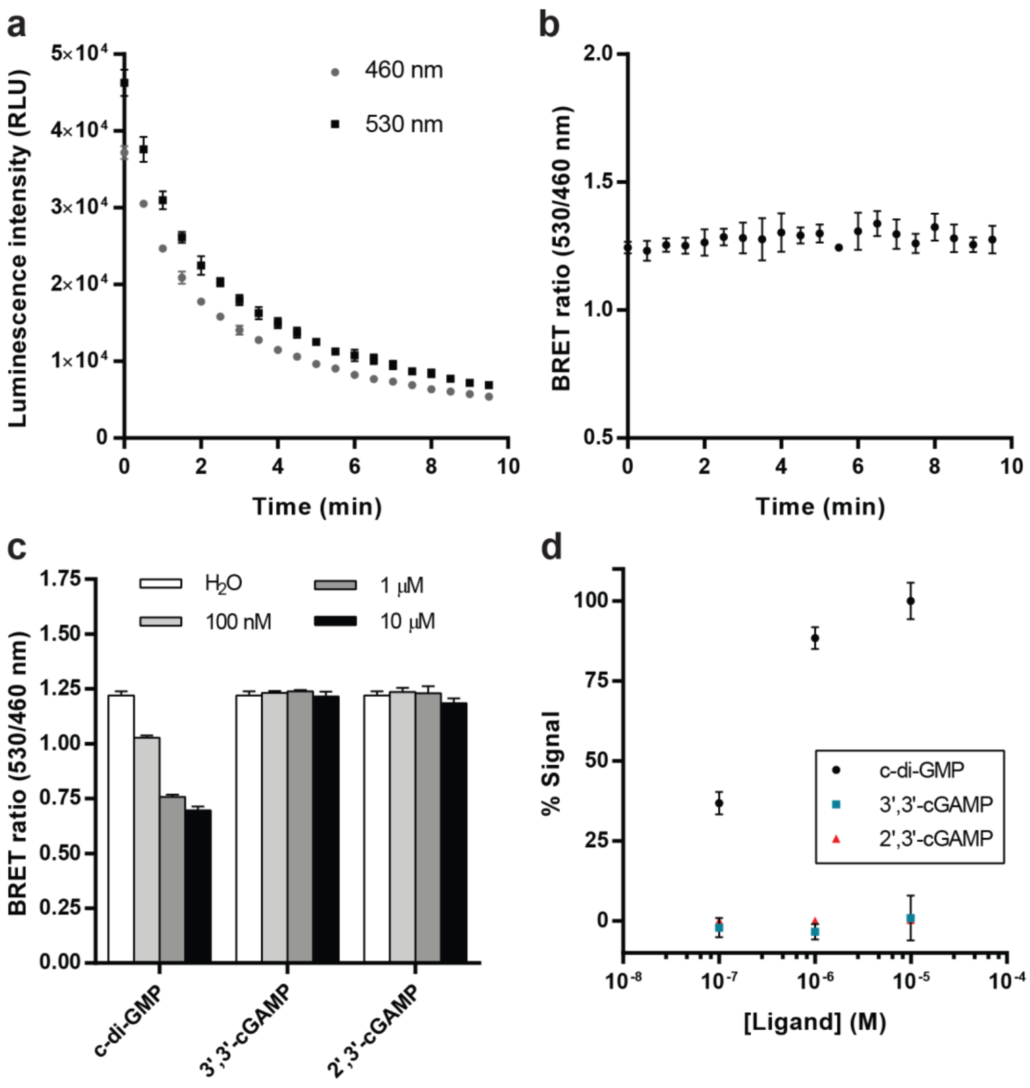


**Figure S1. Additional characterization of VYN-Ec sensor.** (a) Luminescence signal intensity of VYN-Ec sensor over time. Luminescent substrate was added just prior to beginning of measurement. Data are from 3 replicates represented as mean +/- SD. (b) BRET ratio calculated from data shown in part (a). Data are from 3 replicates represented as mean +/- SD. (c) BRET ratios of VYN-Ec sensor in response to different cyclic di-nucleotides. Data are from 3 replicates represented as mean +/- SD. (d) Percent signal calculated from part (c). Data are from 3 replicates represented as mean +/- SD.

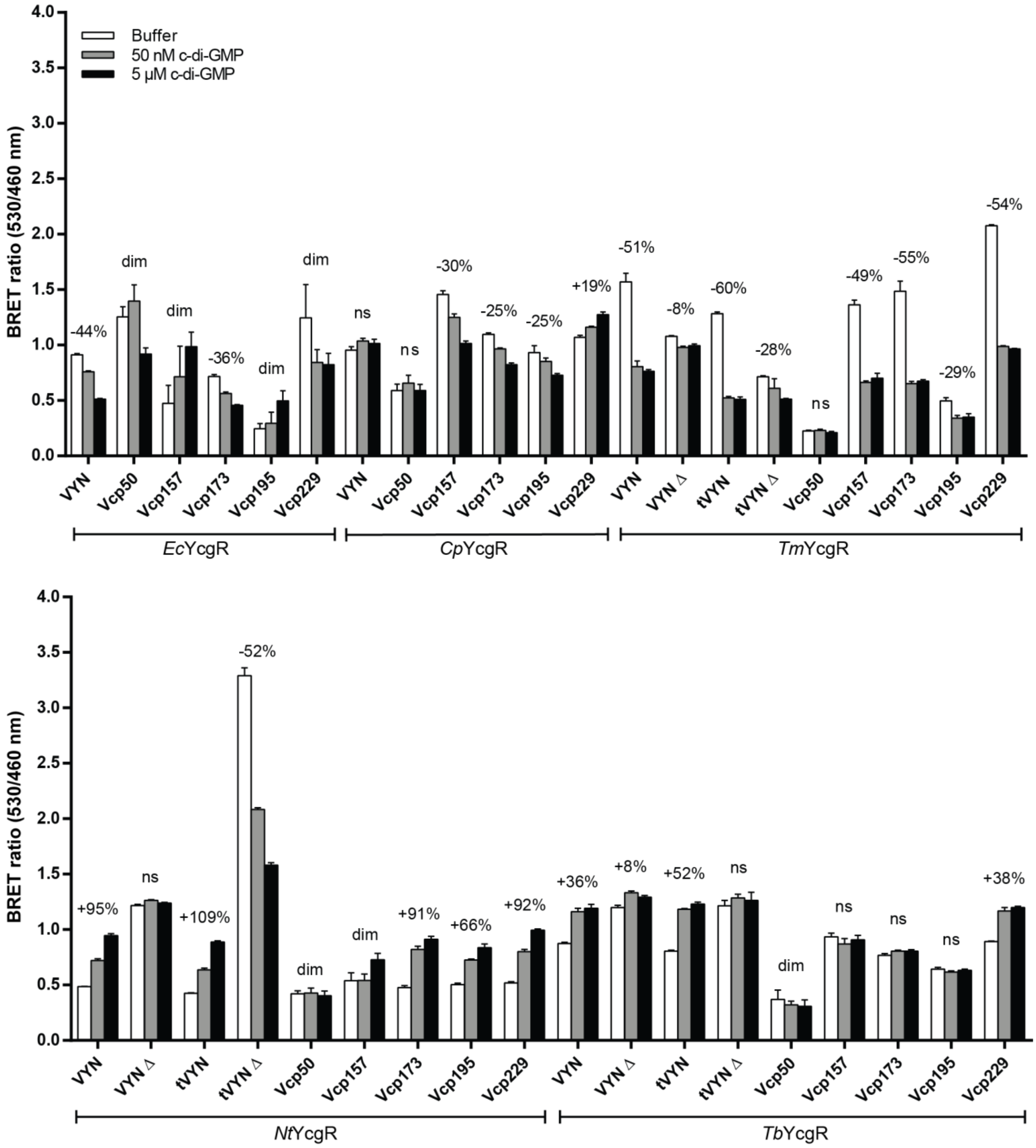


**Figure S2. Lysate-based screen of VYN biosensor library.** BRET value measurements from VYN library lysate screening in response to c-di-GMP (data summarized in Figure 2c). Percent signal change values (comparing buffer to 5 µM c-di-GMP conditions) shown for variants that exhibited significant changes in BRET ($P < 0.05$ determined by Student's t-test). Data are from 4 biological replicates represented as the mean +/- SD.

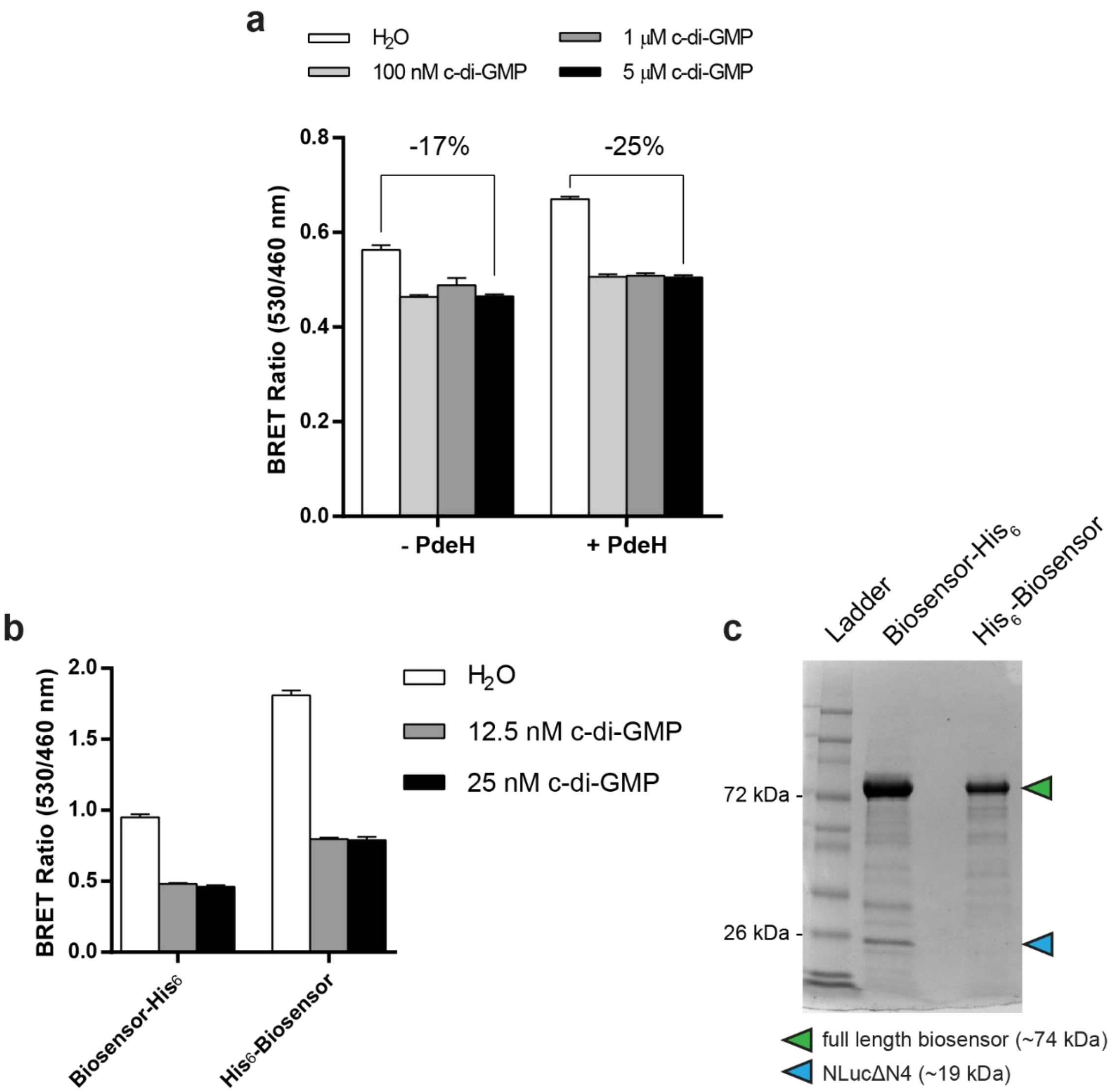


**Figure S3. Additional characterization of tVYN-TmΔ biosensor.** (a) BRET values from the tVYN-TmΔ biosensor in the lysate-based assay with or without the co-expression of the c-di-GMP specific phosphodiesterase, PdeH. Data are from 9 biological replicates represented as the mean +/- SD. (b) BRET values of tVYN-TmΔ *in vitro* after purification with a C-terminal or N-terminal $His_6$ tag. Data are from 3 replicates represented as mean +/- SD. (c) SDS-PAGE of the tVYN-TmΔ biosensor purified with a C-terminal or N-terminal $His_6$ tag, showing the presence of truncated NLuc protein products.

**Differences between biosensor performance in lysates and *in vitro***

For certain sensors, the signal change measured in lysates differs from that measured using purified protein. The most dramatic example of this is for tVYN-TmΔ, which shows a -28% change in the lysate screen and much improved -56% change as purified protein. We investigated this discrepancy and found that the difference likely arises due to two reasons: the extremely high affinity of the sensor, and the generation of luminescent signal from truncated protein products. First, given the extremely high affinity of the sensor, it may be pre-bound to endogenous c-di-GMP present in lysates regardless of the co-expression of PdeH, thereby masking the full signal change. Accordingly, when performing the lysate assay without the co-expression of PdeH, the signal change values are suppressed even further (Figure S3a). Second, we observe different BRET ratios when testing sensor purified with an N-terminal versus a C-terminal affinity tag (Figure S3b). When using a C-terminal affinity tag, truncated protein products containing NLuc are co-purified along with full length biosensor (Figure S3c) These protein products, which are also present in unpurified lysates, produce a high background of “non-specific” luminescent signal at 460 nm. This produces a corresponding decrease in BRET ratio that is independent of the c-di-GMP concentration, partially masking the signal change. Accordingly, when using an N-terminal affinity tag, these truncated protein products are not co-purified, resulting in generally larger BRET ratios and signal change. This finding highlights the fact that for these types of ratiometric sensors, increased “stability” of the scaffold is a highly desirable trait.

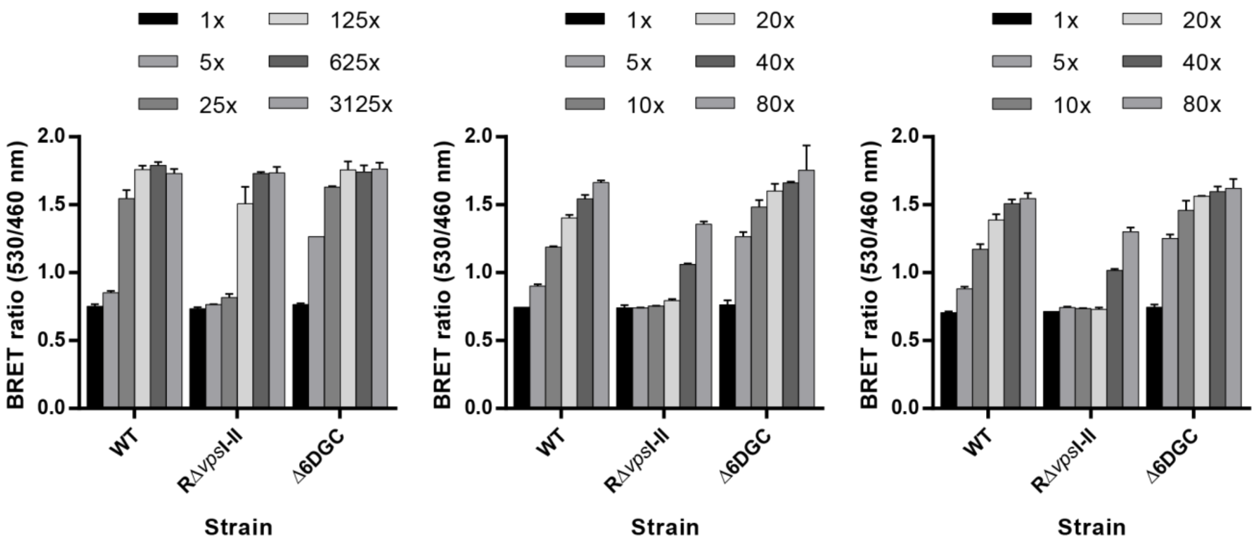


**Figure S4. Biosensor analysis of *V. cholerae* reference strains.** BRET ratios measured in serial dilutions of *V. cholerae* extract, from 1x (no dilution) to 80x or 3125x dilution, using the tVYN-TmΔ biosensor, with each panel representing a separate biological replicate. Note differences in dilution factor for replicate 1 (left) compared to replicates 2 and 3 (middle and right). Data are from 2 technical replicates represented as the mean +/- SD.

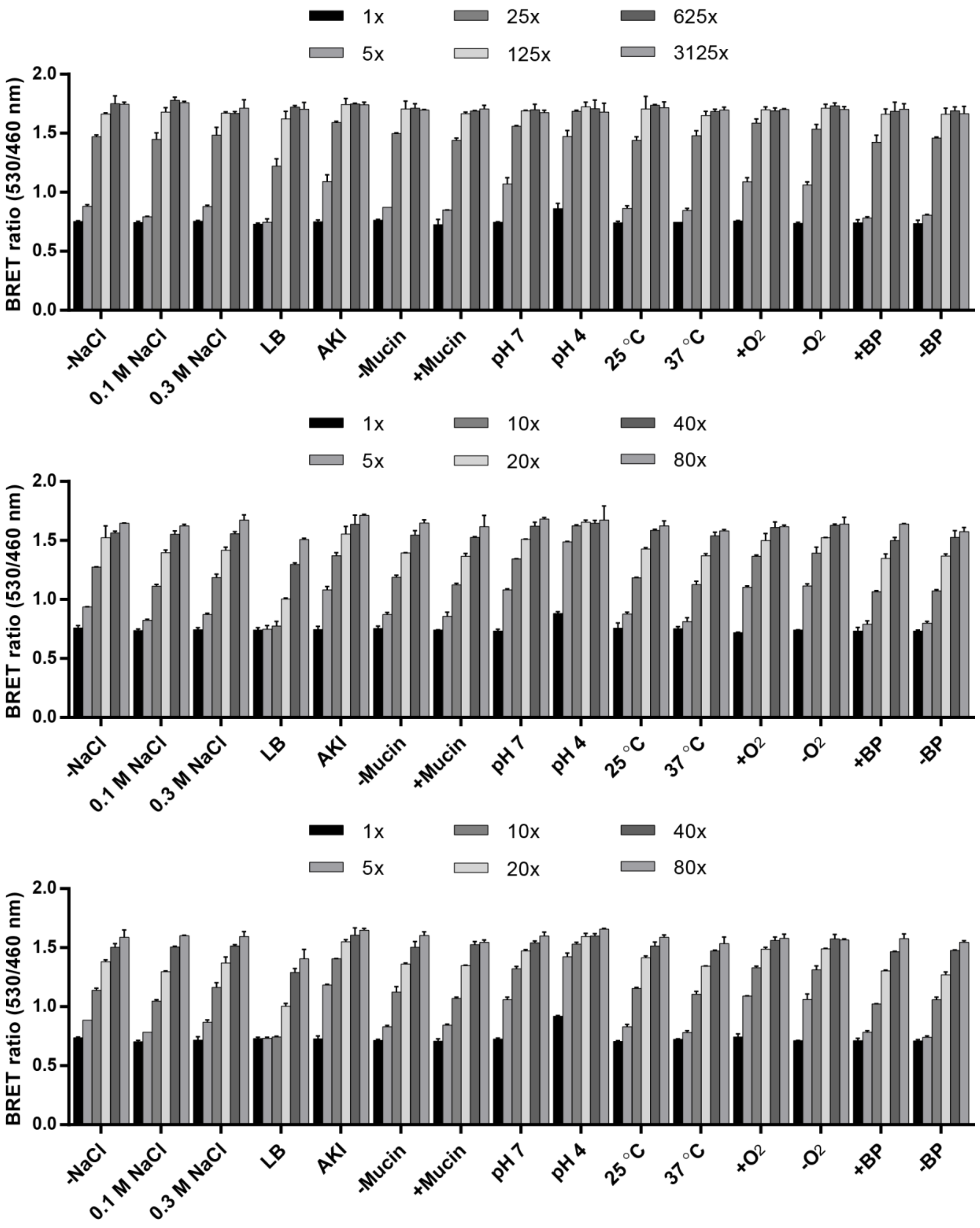


**Figure S5. Biosensor analysis of *V. cholerae* under different growth conditions.** BRET ratios measured in serial dilutions of *V. cholerae* extract, from 1x (no dilution) to 80x or 3125x dilution, using the tVYN-TmΔ biosensor, with each panel representing a separate biological replicate. Note differences in dilution factor for replicate 1 (top) compared to replicates 2 and 3 (bottom). Data are from 2 technical replicates represented as the mean +/- SD.

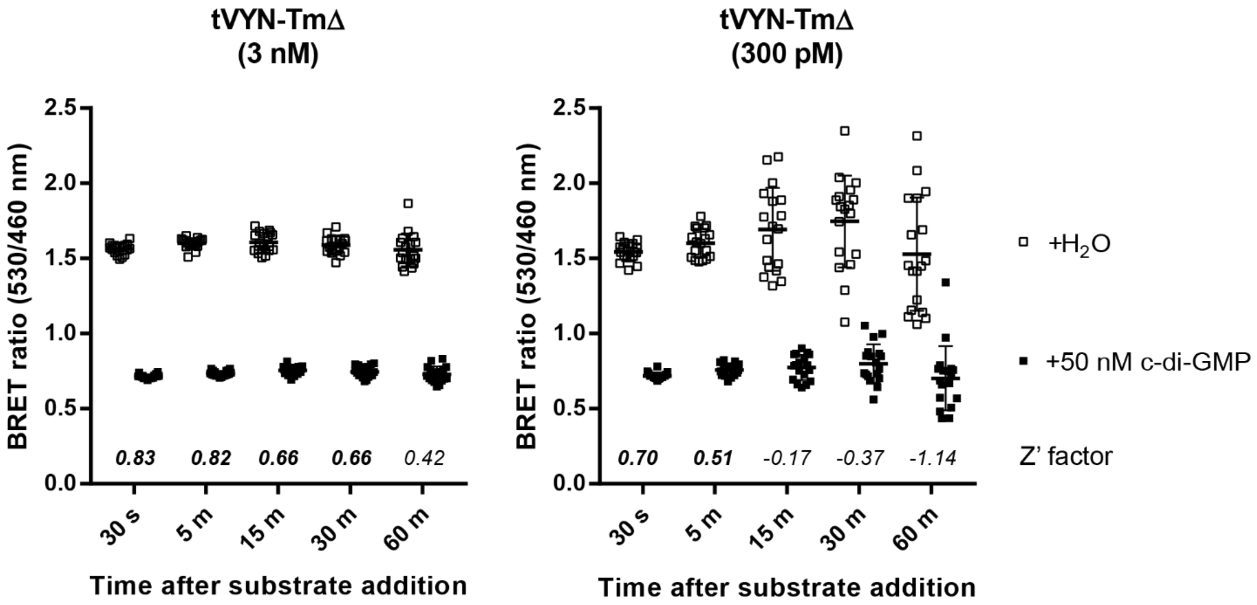


**Figure S6. Determination of Z' factors for plate reader assay.** BRET ratios measured using different concentrations of tVYN-TmΔ for up to 60 minutes after luminescent substrate addition. The calculated Z' factors[4] are reported for each timepoint. Data are from 18 replicates represented as the mean +/- SD.

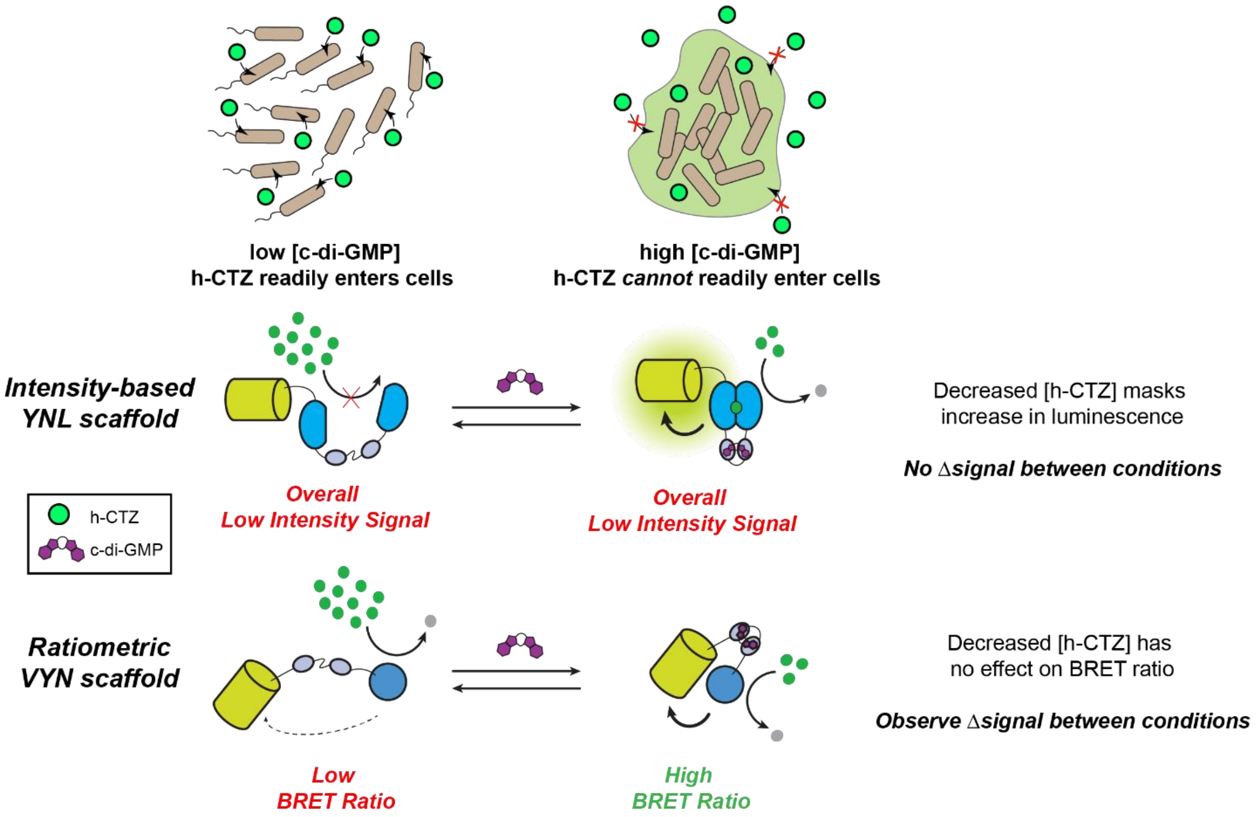


**Figure S7. Live cell imaging with luminescent biosensors.** Schematic showing the difficulties encountered making live cell measurements using the intensity-based YNL scaffold,[2] compared to the ratiometric VYN scaffold.

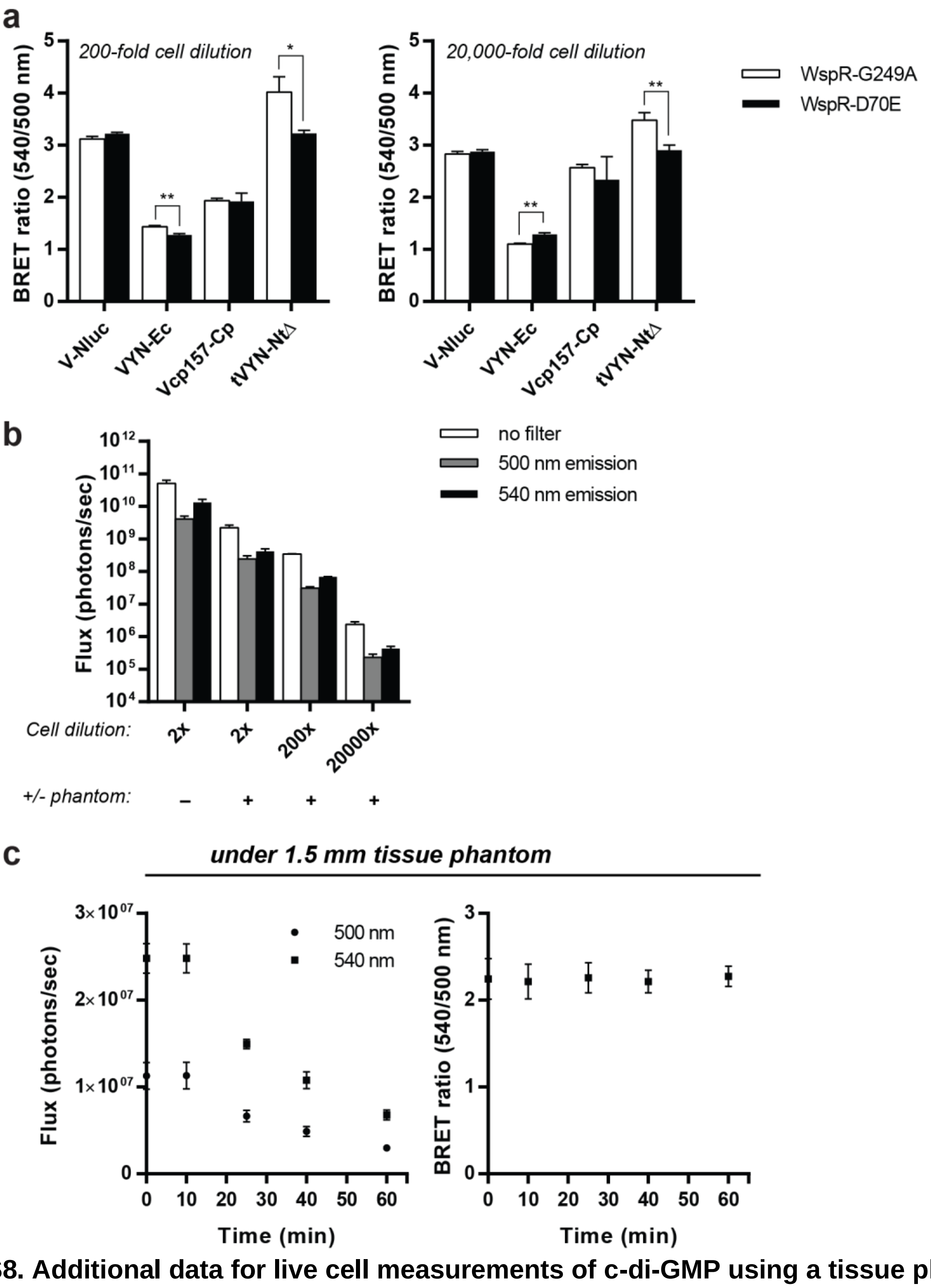


**Figure S8. Additional data for live cell measurements of c-di-GMP using a tissue phantom model.** (a) BRET ratios of serially diluted *E. coli* cultures co-expressing VYN biosensors with WspR-G249A or WspR-D70E calculated from the radiance of each well. Images captured without a phantom cover. Asterisks (*) denote significant changes in BRET ratio (*P < 0.05, **P < 0.005 determined by Student's t-test). (b) Total flux of *E. coli* cultures co-expressing the tVYN-NtΔ biosensor with WspR-G249A at various dilutions with or without the added tissue phantom. (c) (Left) Radiance and (right) corresponding BRET ratios of 200-fold diluted *E. coli* cultures co-expressing the tVYN-NtΔ biosensor with WspR-G249A over time. Luminescent substrate was added at time zero. For all graphs, data are from 3 biological replicates represented as mean +/- SD.

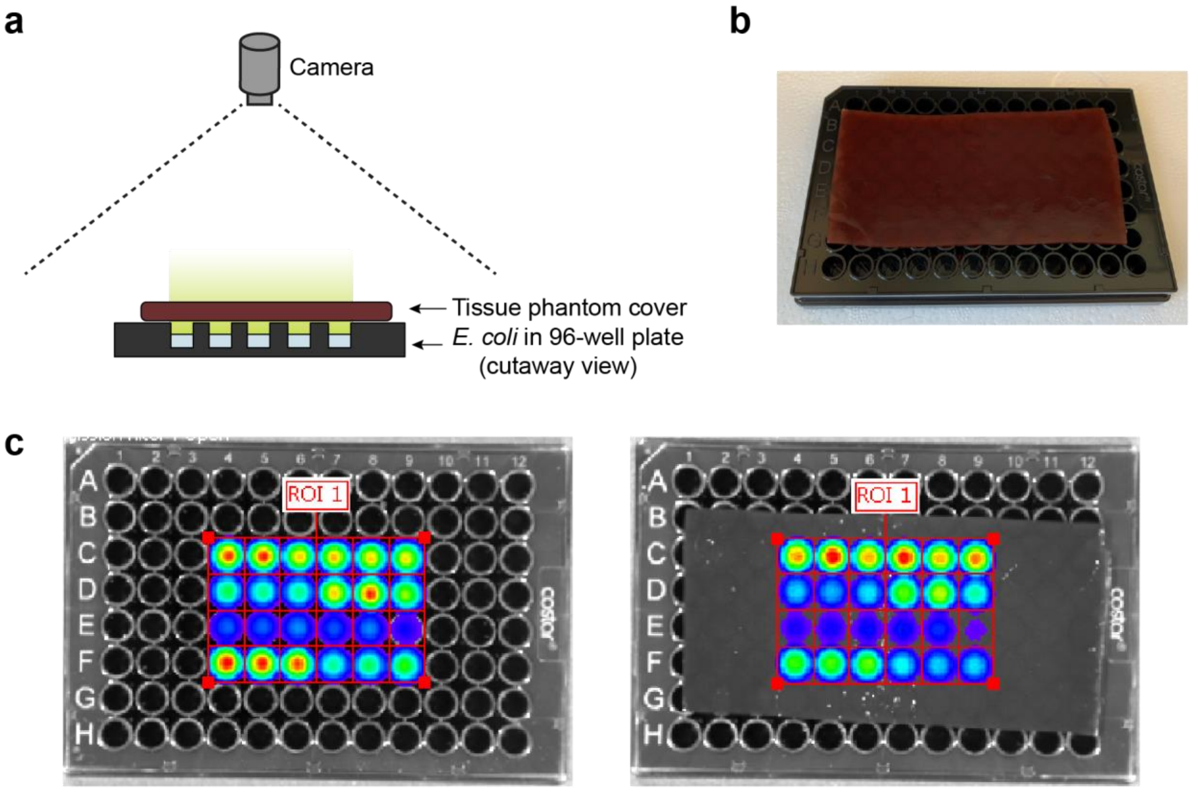


**Figure S9. Tissue phantom imaging setup.** (a) Schematic for imaging with a tissue phantom cover in the IVIS. (b) Representative photograph of plate covered with tissue phantom prior to imaging. (c) Representative images captured by the IVIS showing the ROI grids used for calculating total flux in each well.

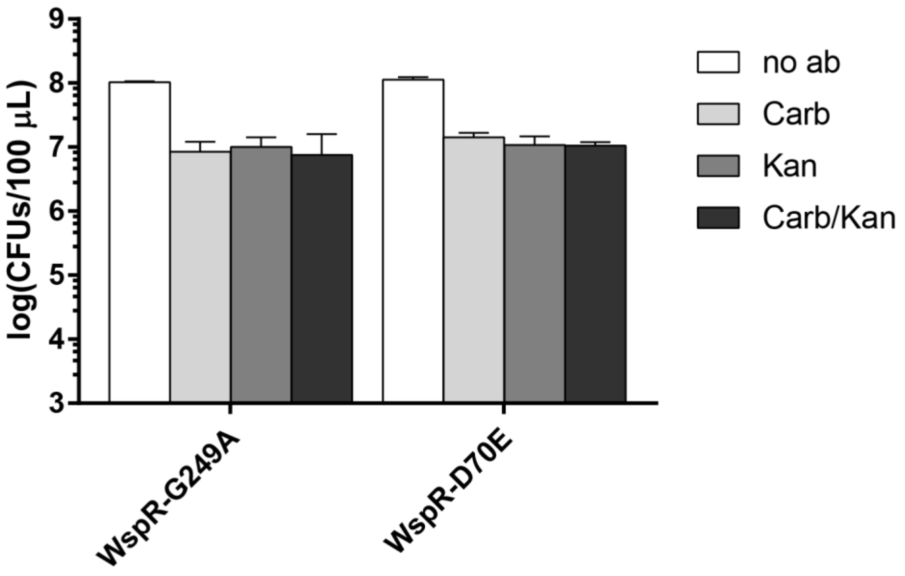


**Figure S10. CFUs in live cell measurements of c-di-GMP.** CFUs measured for *E. coli* co-expressing the tVYN-NtΔ biosensor with WspR-G249A or WspR-D70E after overnight growth in auto-induction media. Data are from 2 biological replicates shown as the mean +/- SD.

**Table S1. Amino acid sequences of biosensor plasmids**

Notes: His-tag, VenusΔC10, NLuc, YcgR; phylogenetic biosensor variants use the same sequences, except the YcgR variant is used in place of TmYcgR (see Table S2); for tVYN and Vcp variants, the corresponding Venus and NLuc sequences were used in place of VenusΔC10 and NLuc (see Table S3)

| | |
|---|---|
| pRSET-VYN | MRGSHHHHHHGMASMTGGQQMGRDLYDDDDKDPMVSKGEELFTGVVPILVELDG DVNGHKFSVSGEGEGDATYGKLTLKLICTTGKLPVPWPTLVTTLGYGLQCFARYPDH MKQHDFFKSAMPEGYVQERTIFFKDDGNYKTRAEVKFEGDTLVNRIELKGIDFKEDG NILGHKLEYNYNSHNVYITADKQKNGIKANFKIRHNIEDGGVQLADHYQQNTPIGDGP VLLPDNHYLSYQSKLSKDPNEKRDHMVLLEFVTAAGGTMVFTLEDFVGDWRQTAGY NLDQVLEQGGVSSLFQNLGVSVTPIQRIVLSGENGLKIDIHVIIPYEGLSGDQMGQIEKI FKVVYPVDDHHFKVILHYGTLVIDGVTPNMIDYFGRPYEGIAVFDGKKITVTGTLWNGN KIIDERLINPDGSLLFRVTINGVTGWRLCERILA*stop |
| pRSET-VYN-Tm | MRGSHHHHHHGMASMTGGQQMGRDLYDDDDKDPMVSKGEELFTGVVPILVELDG DVNGHKFSVSGEGEGDATYGKLTLKLICTTGKLPVPWPTLVTTLGYGLQCFARYPDH MKQHDFFKSAMPEGYVQERTIFFKDDGNYKTRAEVKFEGDTLVNRIELKGIDFKEDG NILGHKLEYNYNSHNVYITADKQKNGIKANFKIRHNIEDGGVQLADHYQQNTPIGDGP VLLPDNHYLSYQSKLSKDPNEKRDHMVLLEFVTAAGGTGMEYYTELVNAKDVIRPGQ NVIVEVSAPEDLEGQYKSSVHDVDFEKRVLTLSMPSFRGRLVPLPRGTRCTVMILDSS AIYVFRTSVLESGRDEDGFPVTKVPFPGRLRKIQRRRFKRIKIFLEGTYRVASRDEPPK RFVTRDFSAGGMLMVVEDILTPEQIIYVTLDLDEDLKLKDHPARVVREAGALETGERM YGVEFLNVPPALERKLVSFVFKKEIEMRNKERSESEGGTMVFTLEDFVGDWRQTAGY NLDQVLEQGGVSSLFQNLGVSVTPIQRIVLSGENGLKIDIHVIIPYEGLSGDQMGQIEKI FKVVYPVDDHHFKVILHYGTLVIDGVTPNMIDYFGRPYEGIAVFDGKKITVTGTLWNGN KIIDERLINPDGSLLFRVTINGVTGWRLCERILA*stop |
| pET21-VYN | MVSKGEELFTGVVPILVELDGDVNGHKFSVSGEGEGDATYGKLTLKLICTTGKLPVP WPTLVTTLGYGLQCFARYPDHMKQHDFFKSAMPEGYVQERTIFFKDDGNYKTRAEV KFEGDTLVNRIELKGIDFKEDGNILGHKLEYNYNSHNVYITADKQKNGIKANFKIRHNIE DGGVQLADHYQQNTPIGDGPVLLPDNHYLSYQSKLSKDPNEKRDHMVLLEFVTAAG GTMVFTLEDFVGDWRQTAGYNLDQVLEQGGVSSLFQNLGVSVTPIQRIVLSGENGLK IDIHVIIPYEGLSGDQMGQIEKIFKVVYPVDDHHFKVILHYGTLVIDGVTPNMIDYFGRPY EGIAVFDGKKITVTGTLWNGNKIIDERLINPDGSLLFRVTINGVTGWRLCERILAKLAAA LEHHHHHH*stop |
| pET21-VYN-Tm | MVSKGEELFTGVVPILVELDGDVNGHKFSVSGEGEGDATYGKLTLKLICTTGKLPVP WPTLVTTLGYGLQCFARYPDHMKQHDFFKSAMPEGYVQERTIFFKDDGNYKTRAEV KFEGDTLVNRIELKGIDFKEDGNILGHKLEYNYNSHNVYITADKQKNGIKANFKIRHNIE DGGVQLADHYQQNTPIGDGPVLLPDNHYLSYQSKLSKDPNEKRDHMVLLEFVTAAG GTGMEYYTELVNAKDVIRPGQNVIVEVSAPEDLEGQYKSSVHDVDFEKRVLTLSMPS FRGRLVPLPRGTRCTVMILDSSAIYVFRTSVLESGRDEDGFPVTKVPFPGRLRKIQRR RFKRIKIFLEGTYRVASRDEPPKRFVTRDFSAGGMLMVVEDILTPEQIIYVTLDLDEDLK LKDHPARVVREAGALETGERMYGVEFLNVPPALERKLVSFVFKKEIEMRNKERSESE GGTMVFTLEDFVGDWRQTAGYNLDQVLEQGGVSSLFQNLGVSVTPIQRIVLSGENG LKIDIHVIIPYEGLSGDQMGQIEKIFKVVYPVDDHHFKVILHYGTLVIDGVTPNMIDYFGR PYEGIAVFDGKKITVTGTLWNGNKIIDERLINPDGSLLFRVTINGVTGWRLCERILAKLA AALEHHHHHH*stop |

**Table S2. Amino acid sequences of YcgR proteins**

Notes: Highlighted residues were removed for YcgRΔ variants

| | |
|---|---|
| EcYcgR | MSHYHEQFLKQNPLAVLGVLRDLHKAAIPLRLSWNGGQLISKLLAITPDKLVLDFGSQ AEDNIAVLKAQHITITAETQGAKVEFTVEQLQQSEYLQLPAFITVPPPTLWFVQRRRYF RISAPLHPPYFCQTKLADNSTLRFRLYDLSLGGMGALLETAKPAELQEGMRFAQIEVN MGQWGVFHFDAQLISISERKVIDGKNETITTPRLSFRFLNVSPTVERQLQRIIFSLEREA REKADKVRD |
| CpYcgR | MAKRKEPKVGDRGILRVREPGETGVEYYSTRIEDVRDGLIACSQPMRGQVYVKILSS PVELTYLKGDSVFSLMCEVIEQGKGDPPLIVLKPISGIYRSDRREYVRVPWMLDAELLF VKTFPADVKKFWEDHSHESVRAVILDLSAGGCRLSLAEACMVGEKVLIRFTVPEPNP DTFLLPALIKRVEPGSEPGVTNVGLQFVDVKDVIRDKLCRSVFCRQRELIKKGFYELEE E |
| TmYcgR | MEYYTELVNAKDVIRPGQNVIVEVSAPEDLEGQYKSSVHDVDFEKRVLTLSMPSFRG RLVPLPRGTRCTVMILDSSAIYVFRTSVLESGRDEDGFPVTKVPFPGRLRKIQRRRFK RIKIFLEGTYRVASRDEPPKRFVTRDFSAGGMLMVVEDILTPEQIIYVTLDLDEDLKLKD HPARVVREAGALETGERMYGVEFLNVPPALERKLVSFVFKKEIEMRNKERSESE |
| NtYcgR | MLKIGLSIKIRVDNKDYSSRIEDMDSDYLYISTPMEKGQLVHFSQGSKISVYIIVKGAVY NFEEKIKEQIKSPVPLLKISKPDKLKKIQRRQFFRLEKKLPVKYKILDDDCESELSDTKD AYALDISGGGLKLATQEIIPVNSFLELNFELNIDEGKNSNIHDIRCVGKIVRTQKVDTDR VSIYHYGVKFISLPSEIQDTIVRFIFNEQRKLRLKGRFSHAKRES |
| TbYcgR | MIKKEELKINQKVEVQIPDGSYKGNYSSRVEEIHPDGSIVLAAPFKRGVLIPLRKGDTVI VNFWGQTAGYSFTTAVLETNYQDVPMIRVAAPSTVRRIQRRNFVRVPAWIPLVFSVS SDSDDPSEKKIYRTETVNVSGGGLLIKSPFKLSEGVCLEMEIHLPKRGPVNARGQVVR VEEKREQSPMYLIGVAFTEIAETDRTKIINFVFEKQREMRQKGLI |

**Table S3. Amino acid sequences for tVYN and Vcp variants**

Notes: For Vcp variants, an additional starting Met and an artificial linker (GGSGG) fusing the original N- and C- termini is added (added start codon, N-terminal portion, C-terminal portion)

| | |
|---|---|
| Vcp50<br>N: (50-239)<br>C: (1-49) | MTTGKLPVPWPTLVTTLGYGLQCFARYPDHMKQHDFFKSAMPEGYVQERTIFFKDD<br>GNYKTRAEVKFEGDTLVNRIELKGIDFKEDGNILGHKLEYNYNSHNVYITADKQKNGIK<br>ANFKIRHNIEDGGVQLADHYQQNTPIGDGPVLLPDNHYLSYQSKLSKDPNEKRDHMV<br>LLEFVTAAGITLGMDELYKGGSGGMVSKGEELFTGVVPILVELDGDVNGHKFSVSGE<br>GEGDATYGKLTLKLIC |
| Vcp157<br>N: (157-239)<br>C: (1-156) | MKQKNGIKANFKIRHNIEDGGVQLADHYQQNTPIGDGPVLLPDNHYLSYQSKLSKDP<br>NEKRDHMVLLEFVTAAGITLGMDELYKGGSGGMVSKGEELFTGVVPILVELDGDVNG<br>HKFSVSGEGEGDATYGKLTLKLICTTGKLPVPWPTLVTTLGYGLQCFARYPDHMKQH<br>DFFKSAMPEGYVQERTIFFKDDGNYKTRAEVKFEGDTLVNRIELKGIDFKEDGNILGH<br>KLEYNYNSHNVYITAD |
| Vcp173<br>N: (173-239)<br>C: (1-172) | MEDGGVQLADHYQQNTPIGDGPVLLPDNHYLSYQSKLSKDPNEKRDHMVLLEFVTA<br>AGITLGMDELYKGGSGGMVSKGEELFTGVVPILVELDGDVNGHKFSVSGEGEGDAT<br>YGKLTLKLICTTGKLPVPWPTLVTTLGYGLQCFARYPDHMKQHDFFKSAMPEGYVQE<br>RTIFFKDDGNYKTRAEVKFEGDTLVNRIELKGIDFKEDGNILGHKLEYNYNSHNVYITA<br>DKQKNGIKANFKIRHNI |
| Vcp195<br>N: (195-239)<br>C: (1-194) | MLLPDNHYLSYQSKLSKDPNEKRDHMVLLEFVTAAGITLGMDELYKGGSGGMVSKG<br>EELFTGVVPILVELDGDVNGHKFSVSGEGEGDATYGKLTLKLICTTGKLPVPWPTLVT<br>TLGYGLQCFARYPDHMKQHDFFKSAMPEGYVQERTIFFKDDGNYKTRAEVKFEGDT<br>LVNRIELKGIDFKEDGNILGHKLEYNYNSHNVYITADKQKNGIKANFKIRHNIEDGGVQL<br>ADHYQQNTPIGDGPV |
| Vcp229<br>N: (229-239)<br>C: (1-228) | MGITLGMDELYKGGSGGMVSKGEELFTGVVPILVELDGDVNGHKFSVSGEGEGDAT<br>YGKLTLKLICTTGKLPVPWPTLVTTLGYGLQCFARYPDHMKQHDFFKSAMPEGYVQE<br>RTIFFKDDGNYKTRAEVKFEGDTLVNRIELKGIDFKEDGNILGHKLEYNYNSHNVYITA<br>DKQKNGIKANFKIRHNIEDGGVQLADHYQQNTPIGDGPVLLPDNHYLSYQSKLSKDPN<br>EKRDHMVLLEFVTAA |
| VenusΔC12 | MVSKGEELFTGVVPILVELDGDVNGHKFSVSGEGEGDATYGKLTLKLICTTGKLPVP<br>WPTLVTTLGYGLQCFARYPDHMKQHDFFKSAMPEGYVQERTIFFKDDGNYKTRAEV<br>KFEGDTLVNRIELKGIDFKEDGNILGHKLEYNYNSHNVYITADKQKNGIKANFKIRHNIE<br>DGGVQLADHYQQNTPIGDGPVLLPDNHYLSYQSKLSKDPNEKRDHMVLLEFVTA |
| NLucΔN4 | LEDFVGDWRQTAGYNLDQVLEQGGVSSLFQNLGVSVTPIQRIVLSGENGLKIDIHVIIP<br>YEGLSGDQMGQIEKIFKVVYPVDDHHFKVILHYGTLVIDGVTPNMIDYFGRPYEGIAVF<br>DGKKITVTGTLWNGNKIIDERLINPDGSLLFRVTINGVTGWRLCERILA |

#### Table S4. Oligonucleotides used in this study

| Name | Sequence |
| --- | --- |
| REV-Venus-Nluc-vector | CCAGTCCCCAACGAAATCTTCGAGTGTGAAGACCATggtaccCCCGGCGG |
| FWD-Venus-Nluc-insert | GTTCGTGACCGCCGCCGGGggtaccATGGTCTTCACACTCGAAGATTTCG |
| FWD-Venus-Nluc-vector | CGGCTGTGCGAACGCATTCTGGCGTAAgAATTCGAAGCTTGATCCGGCTG |
| REV-Venus-Nluc-insert | TTGTTAGCAGCCGGATCAAGCTTCGAATTcTTACGCCAGAATGCGTTCGC |
| REV-VYN-YcgR-vector | ttcaggaactgctcatggtaatgactCATccatggCCCGGCGGCGGTCAC |
| FWD-VYN-YcgR-insert | TCGTGACCGCCGCCGGGccatggATGagtcattaccatgagcagttcctg |
| REV-VYN-YcgR-insert | ACGAAATCTTCGAGTGTGAAGACCATgagctcgtcgcgcactttgtccgc |
| FWD-VYN-YcgR-vector | ggacaaagtgcgcgacgagctcATGGTCTTCACACTCGAAGATTTCGTTG |
| FWD-TmYcgR-VYN-insert | TGACCGCCGCCGGGggtacaggtATGGAATATTATACCGAACTGGTGAAC |
| REV-TmYcgR-VYN-insert | AAATCTTCGAGTGTGAAGACCATggtaccaccTTCGCTTTCGCTGCGTTC |
| FWD-CpYcgR-VYN-insert | GAGTTCGTGACCGCCGCCGGGggtacaggtATGGCGAAACGCAAAGAACC |
| REV-CpYcgR-VYN-insert | CGAGTGTGAAGACCATggtaccaccTTCTTCTTCCAGTTCATAAAAGCCT |
| FWD-TbYcgR-VYN-insert | CGCCGCCGGGggtacaggtATGATTAAAAAAGAAGAACTGAAAATTAACC |
| REV-TbYcgR-VYN-insert | AAATCTTCGAGTGTGAAGACCATggtaccaccAATCAGGCCTTTCTGGCG |
| FWD-VYN-Nt-YcgR-insert | GCCGCCGGGggtacaggtATGCTGAAAATTGGCCTGAGCATTAAAATTCG |
| REV-VYN-Nt-YcgR-insert | GAGTGTGAAGACCATggtaccaccagATTCGCGTTTCGCATGGCTAAAGC |
| FWD-tVYN-rth | CTCGAAGATTTCGTTGGGGACTG |
| REV-tVYN-rth | GGTaccGGCGGTCACGAACTCCA |
| FWD-tVYN-TmYcgR delta flex | CTGCTGGAGTTCGTGACCGCCGGTACaggtTATACCGAACTGGTGAACGC |
| REV-tVYN-TmYcgR delta flex | CCAACGAAATCTTCGAGggtaccaccTTCTTTGTTGCGCATTTCAATTTC |
| FWD-Venus-pET21 insert | TGTTTAACTTTAAGAAGGAGATATACATATGATGGTGAGCAAGGGCGAGG |
| REV-Nluc-pET21 insert | GTGCGGCCGCAAGCTTCGCCAGAATGCGT |
| REV-Nluc-pET21 insert 2 | CGGCCGCAAGCTTCGCCAGAATGCGTTCGC |
| FWD-Tm-Nluc insert | AAGAAGGAGATATACATATGggtacaggtATGGAATATTATACCGAACTG |
| FWD-Venus-pET21 RTH | ATGGACGAGCTGTACAAGAAGCTTGCGGCCGCACTC |
| REV-Venus-pET21 RTH | GCCGAGAGTGATCCCGGCGGCGGTCAC |
| FWD-Tm-Nluc-RTH | ATGGAATATTATACCGAACTGGTGAAC |
| REV-Tm-Nluc-RTH | ATGTATATCTCCTTCTTAAAGTTAAACAAAATTATTTC |
| FWD-cp-linker | GGTGGTTCCGGTGGTATGGTGAGCAAGGGCGAGG |
| REV-cp-linker | ACCACCGGAACCACCCTTGTACAGCTCGTCCATGCC |
| FWD-vcp50 | GTTTAACTTTAAGAAGGAGATATACATATGACCACCGGCAAGCTGCC |

| | |
|---|---|
| REV-vcp50 | CCAGTTCGGTATAATATTCCATacctgtaccGCAGATCAGCTTCAGGGTCAGC |
| FWD-vcp157 | GTTTAACTTTAAGAAGGAGATATACATATGAAGCAGAAGAACGGCATCAAGG |
| REV-vcp157 | CCAGTTCGGTATAATATTCCATacctgtaccGTCGGCGGTGATATAGACGTTG |
| FWD-vcp173 | GTTTAACTTTAAGAAGGAGATATACATATGGAGGACGGCGGCGTG |
| REV-vcp173 | CCAGTTCGGTATAATATTCCATacctgtaccGATGTTGTGGCGGATCTTGAAG |
| FWD-vcp195 | GTTTAACTTTAAGAAGGAGATATACATATGCTGCTGCCCGACAACCAC |
| REV-vcp195 | CCAGTTCGGTATAATATTCCATacctgtaccCACGGGGCCGTCGC |
| FWD-vcp229 | GTTTAACTTTAAGAAGGAGATATACATATGGGGATCACTCTCGGCATGG |
| REV-vcp229 | CCAGTTCGGTATAATATTCCATacctgtaccGGCGGCGGTCACGAACTC |
| REV-Nluc-HisTag | CATAAGCTTCGCCAGAATGCGTTCGCA |
| FWD-vcp-Nluc | AAGCTGATCTGCggtaccATGGTCTTCACACTCGAAGATTTCG |
| REV-vcp50-Nluc | CGAGTGTGAAGACCATggtaccGCAGATCAGCTTCAGGGTCAGC |
| REV-vcp157-Nluc | TGTGAAGACCATggtaccGTCGGCGGTGATATAGACGTTGTGG |
| REV-vcp173-Nluc | GAGTGTGAAGACCATggtaccGATGTTGTGGCGGATCTTGAAG |
| REV-vcp195-Nluc | GAAATCTTCGAGTGTGAAGACCATggtaccCACGGGGCCGTCGC |
| REV-vcp229-Nluc | TTCGAGTGTGAAGACCATggtaccGGCGGCGGTCACGAACTC |
| REV-VYN-Histag | ATCTTCGAGTGTGAAGACCATggtaccCCCGGCGGCGGTCAC |
| FWD-Ec-no vcp | GCCGCCGGGggtacaggtATGagtcattaccatgagcagttc |
| FWD-Ec-vcp50 | CCTGAAGCTGATCTGCggtacaggtATGagtcattaccatgagcagttc |
| FWD-Ec-vcp157 | CTATATCACCGCCGACggtacaggtATGagtcattaccatgagcagttc |
| FWD-Ec-vcp173 | TCCGCCACAACATCggtacaggtATGagtcattaccatgagcagttc |
| FWD-Ec-vcp195 | ACGGCCCGTGggtacaggtATGagtcattaccatgagcagttc |
| FWD-Ec-vcp229 | GACCGCCGCCggtacaggtATGagtcattaccatgagcagttc |
| REV-Ec-vcp | TCGAGTGTGAAGACCATggtaccaccgtcgcgcactttgtccg |
| FWD-Nt-no vcp | GCCGCCGGGggtacaggtATGCTGAAAATTGGCCTGAG |
| FWD-Nt-vcp50 | CCTGAAGCTGATCTGCggtacaggtATGCTGAAAATTGGCCTGAG |
| FWD-Nt-vcp157 | CTATATCACCGCCGACggtacaggtATGCTGAAAATTGGCCTGAG |
| FWD-Nt-vcp173 | TCCGCCACAACATCggtacaggtATGCTGAAAATTGGCCTGAG |
| FWD-Nt-vcp195 | ACGGCCCCGTGggtacaggtATGCTGAAAATTGGCCTGAG |
| FWD-Nt-vcp229 | GACCGCCGCCggtacaggtATGCTGAAAATTGGCCTGAG |
| REV-Nt-vcp | TCGAGTGTGAAGACCATggtaccaccTTCGCGTTTCGCATGG |
| FWD-Tb-no vcp | GCCGCCGGGggtacaggtATGATTAAAAAAGAAGAACTGAAAATTAACC |
| FWD-Tb-vcp50 | TGAAGCTGATCTGCggtacaggtATGATTAAAAAAGAAGAACTGAAAATTAACC |
| FWD-Tb-vcp157 | CACCGCCGACggtacaggtATGATTAAAAAAGAAGAACTGAAAATTAACC |
| FWD-Tb-vcp173 | CCACAACATCggtacaggtATGATTAAAAAAGAAGAACTGAAAATTAACC |
| FWD-Tb-vcp195 | CGGCCCCGTGggtacaggtATGATTAAAAAAGAAGAACTGAAAATTAACC |

| | |
|---|---|
| FWD-Tb-vcp229 | GACCGCCGCCggtacaggtATGATTAAAAAAGAAGAACTGAAAATTAACC |
| REV-Tb-vcp | TCGAGTGTGAAGACCATggtaccaccAATCAGGCCTTTCTGGCG |
| FWD-Cp-no vcp | GCCGCCGGGggtacaggtATGGCGAAACGCAAAGAAC |
| FWD-Cp-vcp50 | CCTGAAGCTGATCTGCggtacaggtATGGCGAAACGCAAAGAAC |
| FWD-Cp-vcp157 | CTATATCACCGCCGACggtacaggtATGGCGAAACGCAAAGAAC |
| FWD-Cp-vcp173 | TCCGCCACAACATCggtacaggtATGGCGAAACGCAAAGAAC |
| FWD-Cp-vcp195 | ACGGCCCCGTGggtacaggtATGGCGAAACGCAAAGAAC |
| FWD-Cp-vcp229 | GACCGCCGCCggtacaggtATGGCGAAACGCAAAGAAC |
| REV-Cp-vcp | TCGAGTGTGAAGACCATggtaccaccTTCTTCTTCCAGTTCATAAAAGCC |
| FWD-VYN-His tag | TGTTTAACTTTAAGAAGGAGATATACATATGGTGAGCAAGGGCGAG |
| REV-VYN-His tag 2 | GTGCGGCCGCAAGCTTCGCCAGAATGCGTTCGC |
| REV-Nt-VYN insert | GTGAAGACCATggtaccaccAGATTCGCGTTTCGCATG |
| FWD-tVYN Nt-insert | GACCGCCGGTACaggtATGCTGAAAATTGGCCTGAG |
| REV-tVYN Nt-insert | CGAAATCTTCGAGggtaccaccAGATTCGCGTTTCGCATG |
| FWD-tVYN Tb-insert | GTGACCGCCGGTACaggtATGATTAAAAAAGAAGAACTGAAAATTAACC |
| REV-tVYN Tb-insert | CAACGAAATCTTCGAGggtaccaccAATCAGGCCTTTCTGGCG |
| FWD-Nt delta-VYN insert | CCGCCGGGggtacaggtAGCATTAAAATTCGCGTGG |
| REV-Nt delta-VYN insert | TCGAGTGTGAAGACCATggtaccaccCAGGCGCAGTTTGCG |
| FWD-Tb delta-VYN insert | GCCGCCGGGggtacaggtAAAGTGGAAGTGCAGATTCCG |
| REV-Tb delta-VYN insert | TCGAGTGTGAAGACCATggtaccaccCTGGCGCATTTCGCG |
| FWD-Tm delta-vcp229 | GACCGCCGCCggtacaggtTATACCGAACTGGTGAACGCG |
| REV-Tm delta-vcp229 | GAGTGTGAAGACCATggtaccaccTTCTTTGTTGCGCATTTCAATTTC |
| FWD-Nt delta-vcp229 | GACCGCCGCCggtacaggtAGCATTAAAATTCGCGTGGATAAC |
| REV-Nt delta-vcp229 | CGAGTGTGAAGACCATggtaccaccCAGGCGCAGTTTGCG |
| FWD-Tb delta-vcp229 | GACCGCCGCCggtacaggtAAAGTGGAAGTGCAGATTCCG |
| REV-Tb delta-vcp229 | CGAGTGTGAAGACCATggtaccaccCTGGCGCATTTCGCG |
| FWD-Nt delta-vcp173 | CCGCCACAACATCggtacaggtAGCATTAAAATTCGCGTGGATAAC |
| FWD-Nt delta-tVYN insert | CGTGACCGCCGGTACaggtAGCATTAAAATTCGCGTGG |
| REV-Nt delta-tVYN insert | CCAACGAAATCTTCGAGggtaccaccCAGGCGCAGTTTGCG |
| FWD-Tb delta-tVYN insert | TCGTGACCGCCGGTACaggtAAAGTGGAAGTGCAGATTCCG |
| REV-Tb delta-tVYN insert | CAACGAAATCTTCGAGggtaccaccCTGGCGCATTTCGCG |
| FWD-Tm delta-tVYN insert | TCGTGACCGCCGGTACaggtTATACCGAACTGGTGAACGC |
| REV-Tm delta-tVYN insert | CAACGAAATCTTCGAGggtaccaccTTCTTTGTTGCGCATTTCAATTTC |
| FWD-tVYN Tm-insert | TTCGTGACCGCCGGTACaggtATGGAATATTATACCGAACTGGTGAAC |
| REV-tVYN Tm-insert | CAACGAAATCTTCGAGggtaccaccTTCGCTTTCGCTGCGTTC |
| FWD-Tm delta-VYN insert | GCCGCCGGGggtacaggtTATACCGAACTGGTGAACGCG |
| REV-Tm delta-VYN insert | TCGAGTGTGAAGACCATggtaccaccTTCTTTGTTGCGCATTTCAATTTC |
